# Learning-dependent evolution of odor mixture representations in piriform cortex

**DOI:** 10.1101/2022.08.25.505306

**Authors:** Alice Berners-Lee, Elizabeth Shtrahman, Julien Grimaud, Venkatesh N. Murthy

**Affiliations:** Department of Molecular and Cellular Biology and Center for Brain Science, Harvard University, Cambridge, MA, USA; Cell Engineering Laboratory (CellTechs), Sup’Biotech, Villejuif, France

## Abstract

Rodents can learn from exposure to rewarding odors to make better and quicker decisions. The piriform cortex is thought to be important for learning complex odor associations, however it is not understood exactly how it learns to remember discriminations between many, sometimes overlapping, odor mixtures. We investigated how odor mixtures are represented in the posterior piriform cortex (pPC) of mice while they learn to discriminate a unique target odor mixture against hundreds of nontarget mixtures. We find that a significant proportion of pPC neurons discriminate between the target and all other nontarget odor mixtures. Neurons that prefer the target odor mixture tend to respond with brief increases in firing rate at odor onset compared to other neurons, which exhibit sustained and/or decreased firing. We allowed mice to continue training after they had reached high levels of performance and find that pPC neurons become more selective for target odor mixtures as well as for randomly chosen repeated nontarget odor mixtures that mice did not have to discriminate from other nontargets. These single unit changes during overtraining are accompanied by better categorization decoding at the population level, even though behavioral metrics of mice such as reward rate and latency to respond do not change. However, when difficult ambiguous trial types are introduced, the robustness of the target selectivity is correlated with better performance on the difficult trials. Taken together, these data reveal pPC as a dynamic and robust system that can optimize for both current and possible future task demands at once.

## Introduction

Rodents depend on odor learning for many important behaviors (Doty, 1986; Raithel and Gottfried, 2021). The piriform cortex, a key olfactory cortical region, clearly reports single odors with passive exposure, with about 10-20% of piriform neurons encoding a single odor (Bolding and Franks, 2017; Poo and Isaacson, 2009; Roland et al., 2017; Stettler and Axel, 2009). The piriform cortex is thought to be important in associative learning (Blazing and Franks, 2020; Gottfried, 2010; Haberly and Bower, 1989; Wilson and Sullivan, 2011). Indeed, when a single odor is paired with reward, although piriform neurons do not encode odor valence (Gadziola et al., 2020; Millman and Murthy, 2020), more piriform neurons are often recruited to respond to those odors compared to unrewarded odors (Gire et al., 2013; Roesch et al., 2006). In the posterior piriform cortex (pPC) specifically, more neurons are recruited to distinguish between two single odors that can lead to obtaining reward or avoiding punishment (Calu et al., 2007).

However, in a rodent’s natural environment odors can be more complex and there may be hundreds of overlapping odors to need to distinguish from a goal or target odor. Mice can distinguish a single odor within a large odor mixture (Rokni et al., 2014), but in that case they have access to all relevant odors and do not have to depend on memory. It is unclear how reward-learning changes pPC encoding in situations where mice may be exposed to more mixtures, one at a time, and how these representations changes with experience.

We developed a complex odor task where mice learned to categorize one target odor mixture differently than all the hundreds of other odor mixtures. After a quick learning phase, we recorded from pPC neurons. We describe how pPC neurons encode odor cues in the task, and then observe how that representation changes with overlearning. We observed that the pPC over-represents important odor mixtures and continues to do so even without reaping any immediate benefit in reward rate. Finally, we introduce “probe” trials that challenge the mouse with ambiguous odor-mixtures and observe how the representation in pPC leverages its robust coding to overcome the novel challenge trials. Taken together, our data reveal a robust and constantly dynamic pPC, that can flexibly prepare for the possible future.

## Results

### Mice learn to discriminate a unique target odor mixture

Four head-fixed water-restricted male mice were presented with an odor mixture made up of three odors (Figure 1). Mice were trained to lick left for the unique target odor mixture (Ethyl tiglate, Allyl tiglate, and Methyl tiglate) and right for any other combination of odors (Figure 1ab). In each trial, the odor was presented for two seconds and a lick to either port in the 2500 ms after odor onset was counted as a response and rewarded for correct responses (Figure 1d). Mice learned the task within a few days (“Learning”; Figure 1e; Table 1, row 2). In separate cohorts of mice, other target odor mixtures were used, and mice were similarly adept at performing the task, arguing for the generality of our results (Supplementary Figure 1). After this initial learning, mice continued to perform the task (“Overlearning”; Figure 1f; days 8-28; Table 1, row 3). In order to compare neural responses to learned (target) and unlearned mixtures, during these sessions some of the nontarget trials were repeated multiple times (“nontarget repeats”; Figure 1b). In later sessions, probe mixtures were introduced (Figure 1c,g; days 22-38; Table 1, row 4), where a mixture contained one of the three odors from the target mixture along with two randomly chosen nontarget odors. Mice were rewarded for classifying both probe and nontarget repeat trials as nontarget mixtures (using the criterion that all 3 components must be present for it to be a target odor). Mice improved their accuracy in nontarget and target trials across sessions only in the learning stage of the task, not in later stages where they appear to be consistently at ceiling (significant interaction between stage and session, Table 1, row 1; session is significant in the learning and probe stages but not the overlearning stage, Table 1, rows 2-4). These data show that mice can quickly learn to tell apart a target odor mixture from hundreds of other non-target mixtures, and further learn to place ambiguous odor mixtures in the correct category.

**Figure 1.**
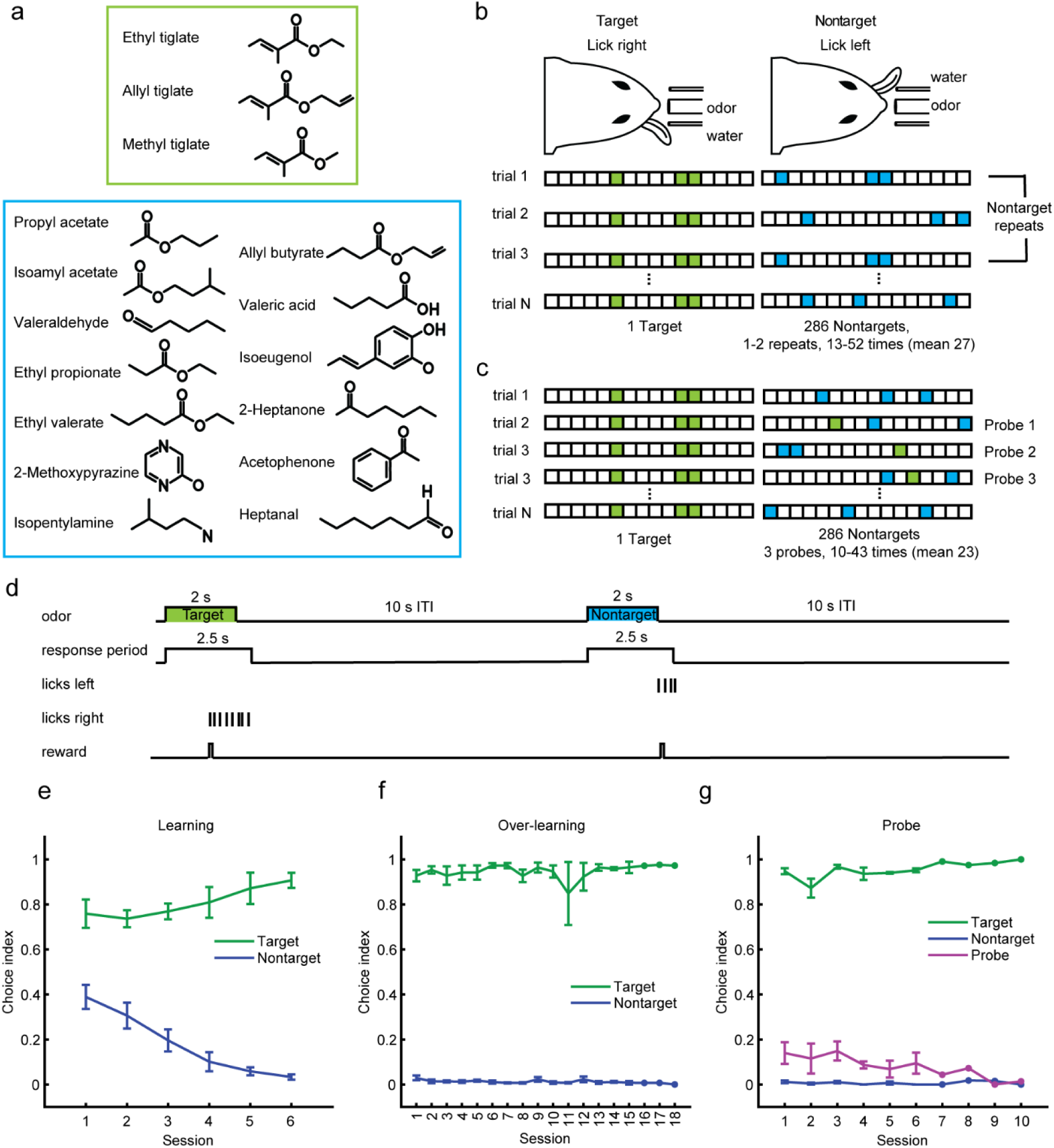
Task and behavior. **(a)** Odor panel. Three odor components make up the target mixture (top) and the nontarget odors are made up of 13 other components (bottom). **(b)** Behavioral set up (top) and stimulus structure (bottom). Colored boxes mark the odorants present in each trial (green for target, blue for nontarget). Some nontarget trials were repeated while others only appeared once. **(c)** Probe component mixture task. Some of the nontarget trials are probe component mixtures containing one target odorant and two nontarget odorants. Mice receive water rewards for categorizing probe mixtures as nontargets (left lick choice). **(d)** Trial structure. Example trial structure for a correct target and nontarget trial. There was a two second odor duration followed by a ten second intertrial-interval (ITI). The response period was the entirety of the odor duration and the following 500 ms. **(e-g)** Target/nontarget task performance of four mice for multiple sessions. Choice index is 1 for lick right, and 0 for lick left. Sessions arranged in order mice performed them. Mice change their choices for target and nontarget trials across sessions during the Learning stage but not following two stages (see Table 1, rows 1-4). Error bars represent mean +/− S.E.M. across mice. **(e)** Learning phase, with only target and nontarget trials (no electrophysiology recordings during this time). Session 1 for each mouse is the first session with blocked trials (see methods for more details on training). **(f)** Overlearning. Session 1 is the first session with random trial structure. **(g)** Sessions with probe trials. Session 1 is when probes were first introduced.

**Table 1:**
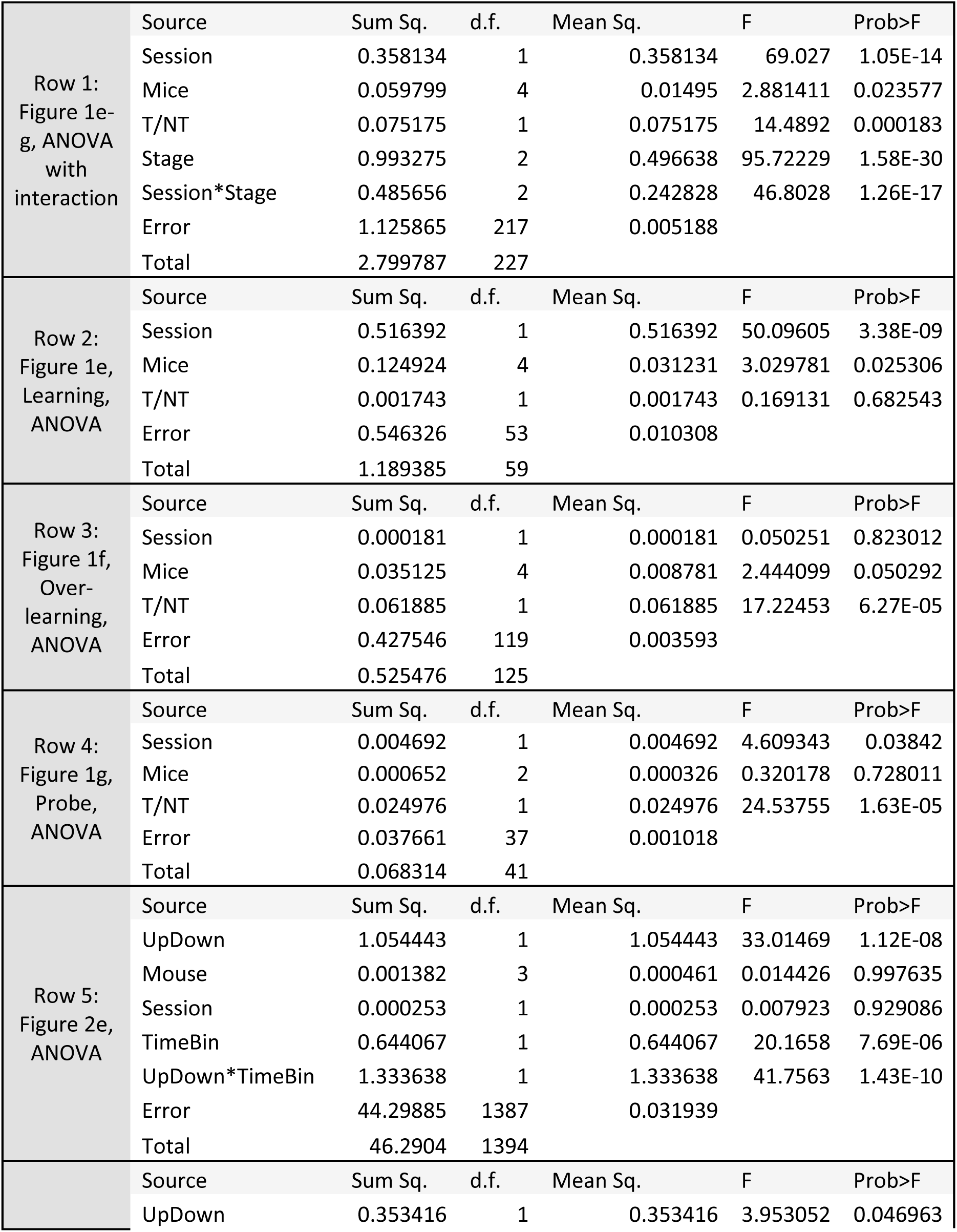
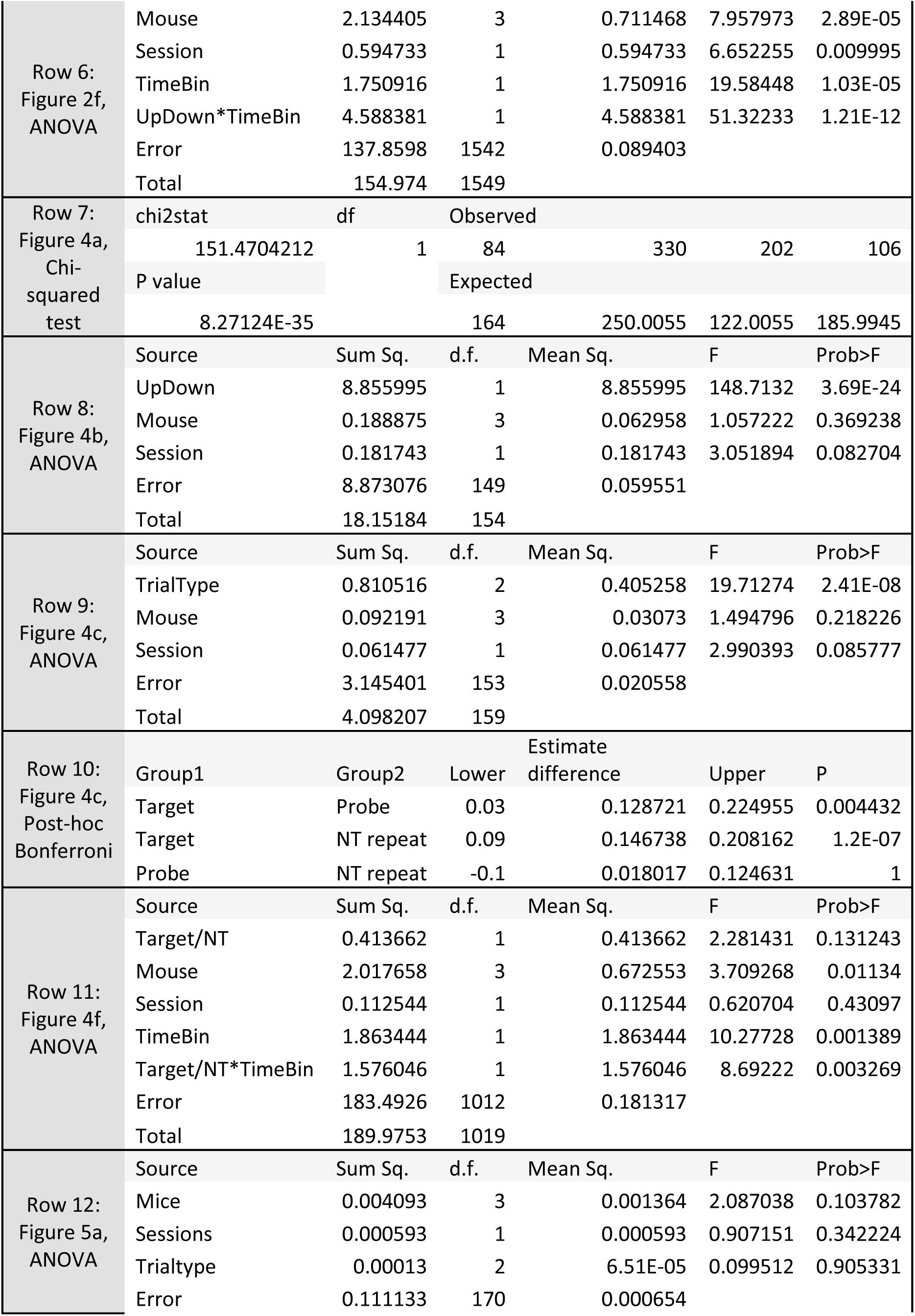
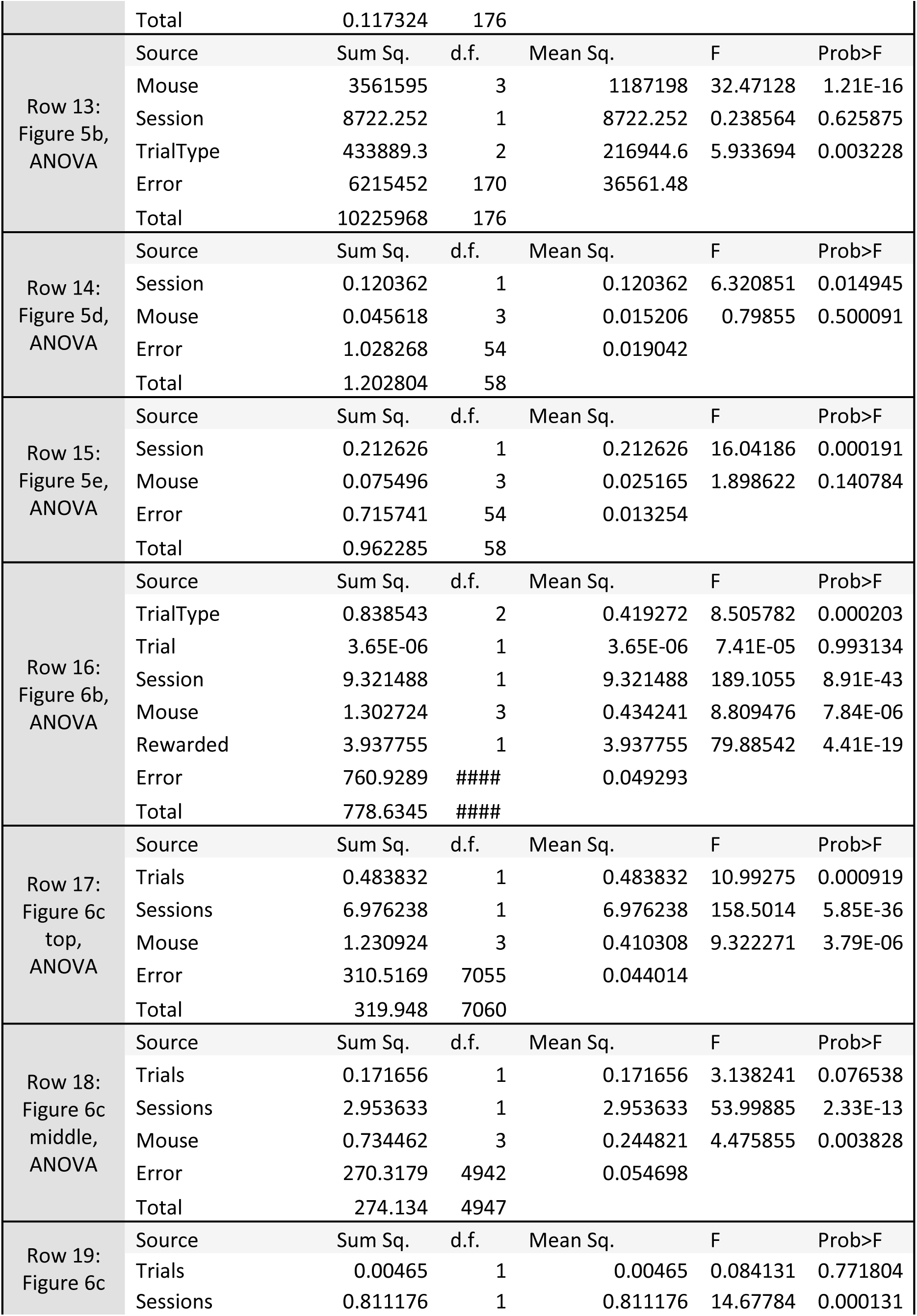
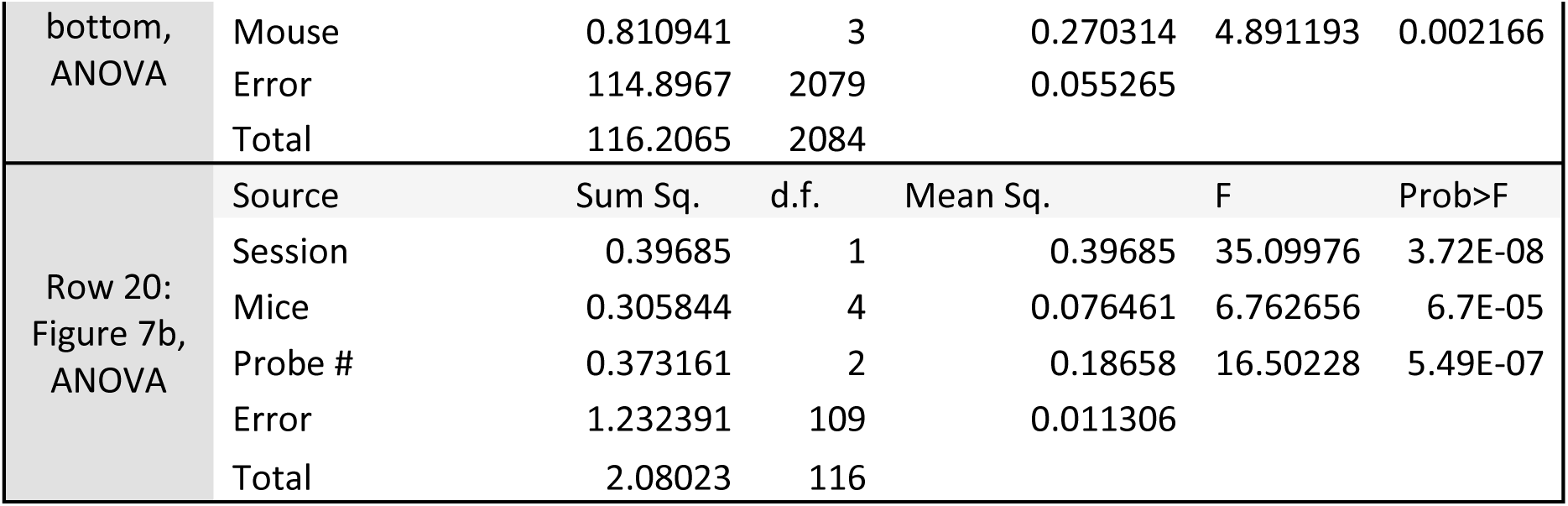
Statistics of comparisons.

### Piriform neurons show brief upward or prolonged downward modulations at odor onset

Tetrodes were implanted in the posterior piriform cortex (0.5 mm posterior and 3.8 mm lateral from bregma, and 3.8 mm ventral from the brain surface; Supplementary Figure 2) and spikes were clustered to isolate single putative pyramidal neurons (Supplementary Figure 3). We found that the majority of neurons are modulated at odor onset (Figure 2). We also found that some neurons respond to the first lick the mouse made (Supplementary Figure 4; neurons with such responses were excluded from counting as odor onset modulated neurons) as well as a small proportion that respond to odor offset (Supplementary Figure 5). We focused subsequent analyses on the largest portion of task-relevant neurons, those modulated at odor onset.

**Figure 2.**
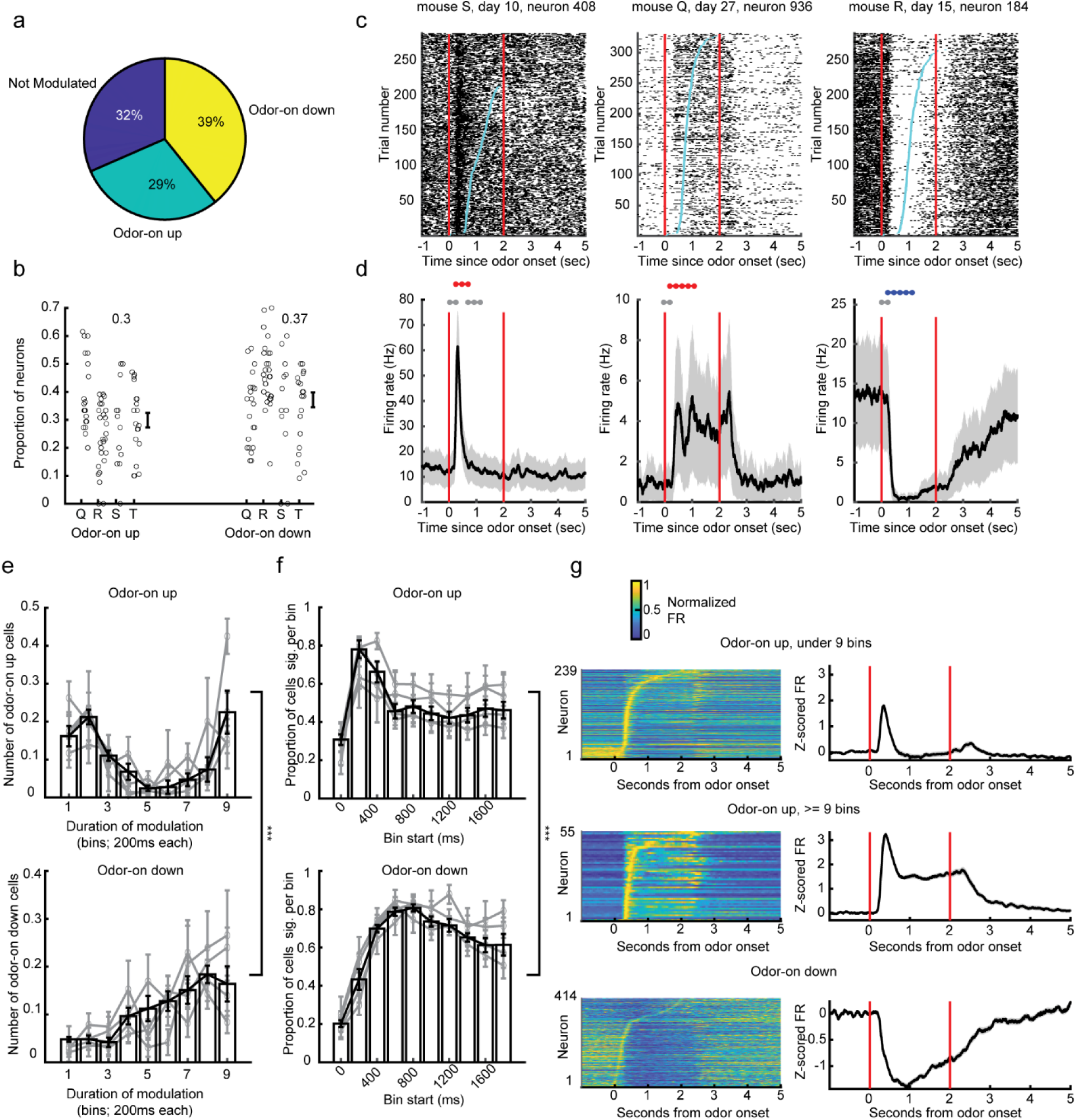
Piriform neurons respond to odor onset. **(a)** Proportion of all piriform neurons that significantly respond to odor onset by either increasing (odor-on up; green) or decreasing (odor-on down; yellow) their firing rate. **(b)** Proportion of neurons that significantly increase or decrease their firing rate across all sessions for all four mice (each small circle is one session). The mean proportion is in text above each condition. **(c)** Raster plots of three example neurons that respond positively (left two) and negatively (right) to odor onset. Red lines depict odor onset and offset, light blue dots depict lick times. **(d)** Peri-stimulus time histograms (PSTHs) of the same three example neurons as in **(c)**. Time bins tested for significant modulation are marked above each trace (red is significantly positive, blue significantly negative). P < 0.05 adjusted for multiple comparisons. **(e, f)** The number of time bins in which neurons are significantly modulated (left) and the proportion of neurons that are significantly modulated in each time bin (right) are plotted for neurons that are significantly upwardly modulated at odor onset (odor-on up, panel e) and significantly downwardly modulated at odor onset (odor-on down, panel f). Gray lines represent the mean and S.E.M for each mouse separately. ANOVA details in Table 1, rows 5 and 6. **(g)** The normalized firing rate (left) and baseline-subtracted z-scored firing rates (right) for upwardly modulated neurons with fewer than 9 200 ms time bins significantly upwardly modulated (top), at least 9 200 ms time bins significantly upwardly modulated (middle) and downwardly modulated neurons (bottom). Red lines depict odor onset and offset. All error bars and shading represent mean +/− S.E.M. *** P < 1e-11.

Across all sessions, 308 of 1055 neurons isolated from tetrode recordings (Figure 2a), an average of 29% of neurons a session (Figure 2b), have significantly higher firing in the second after odor onset (Figure 2c-d, left two examples) compared to baseline (see methods). In addition, 414 of 1055 neurons (Figure 2a), an average of 39% of neurons a session (Figure 2b), have significantly reduced firing in the second after odor onset (Figure 2c-d, right example) compared to baseline (see methods).

We measured the total duration that neurons remain upwardly modulated after odor onset (“Odor-on up”), and found the distribution of these durations to be bimodal (Figure 2e, top); many neurons responded with brief modulations, and another set responded with prolonged modulations (Figure 2g, top and middle), most frequently modulated within 200-600ms after odor onset (Figure 2g, top). By contrast, neurons that reduce their firing rates after odor onset (“Odor-on down”) tend to have prolonged modulations (Figure 2e, bottom) across most of the odor period (Figure 2f-g, bottom). Whether we consider the duration of each neuron’s modulation (Figure 2e; Table 1, row 5) or the proportion of neurons modulated per time-bin (Figure 2f; Table 1, row 6), the odor-on up neurons tend to respond earlier and more briefly than odor-on down neurons.

Taken together, these data show that most piriform neurons are modulated at odor onset, and that the direction and time course of their modulation are related.

### Piriform neurons respond differently to target and nontarget odor mixtures

We observed many neurons that have a higher firing rate for nontarget odor (Figure 3a; nontarget preferring) as well as those that have a higher firing rate during the target odor (Figure 3b; target preferring). There is a significant portion of odor-on upwardly modulated and downwardly modulated neurons that significantly differentiate between target and nontarget odor mixtures, but odor-on up neurons constitute a significantly greater proportion (Figure 4a-b; Table 1, rows 7-8). To test how unique this large proportion of target selective neurons was, we compared it to the proportion of neurons that were selective for probe or nontarget repeat odor mixtures. Nontarget repeats, which are randomly chosen from mixture of nontarget odors that mice do not need to differentiate from other nontargets, are particularly useful as comparisons to the learning of specific target mixture. We found that the target/nontarget discrimination is represented by significantly more neurons than either probe or nontarget repeat trial types (Figure 4c; Table 1, rows 9-10). This is still the case when we down-sampled the number of trials to match between conditions and made a paired comparison across sessions (Supplementary Figure 6). We explored the dynamics of the target-preferring (Figure 4d-f left) to the nontarget preferring (Figure 4d-f right) neurons and noticed that the target preferring neurons tend to be modulated early after odor onset (especially 200-600 ms bins), whereas the nontarget preferring neurons have a greater likelihood of being modulated in later time-bins as well (Figure 4f; Table 1, row 11).

**Figure 3.**
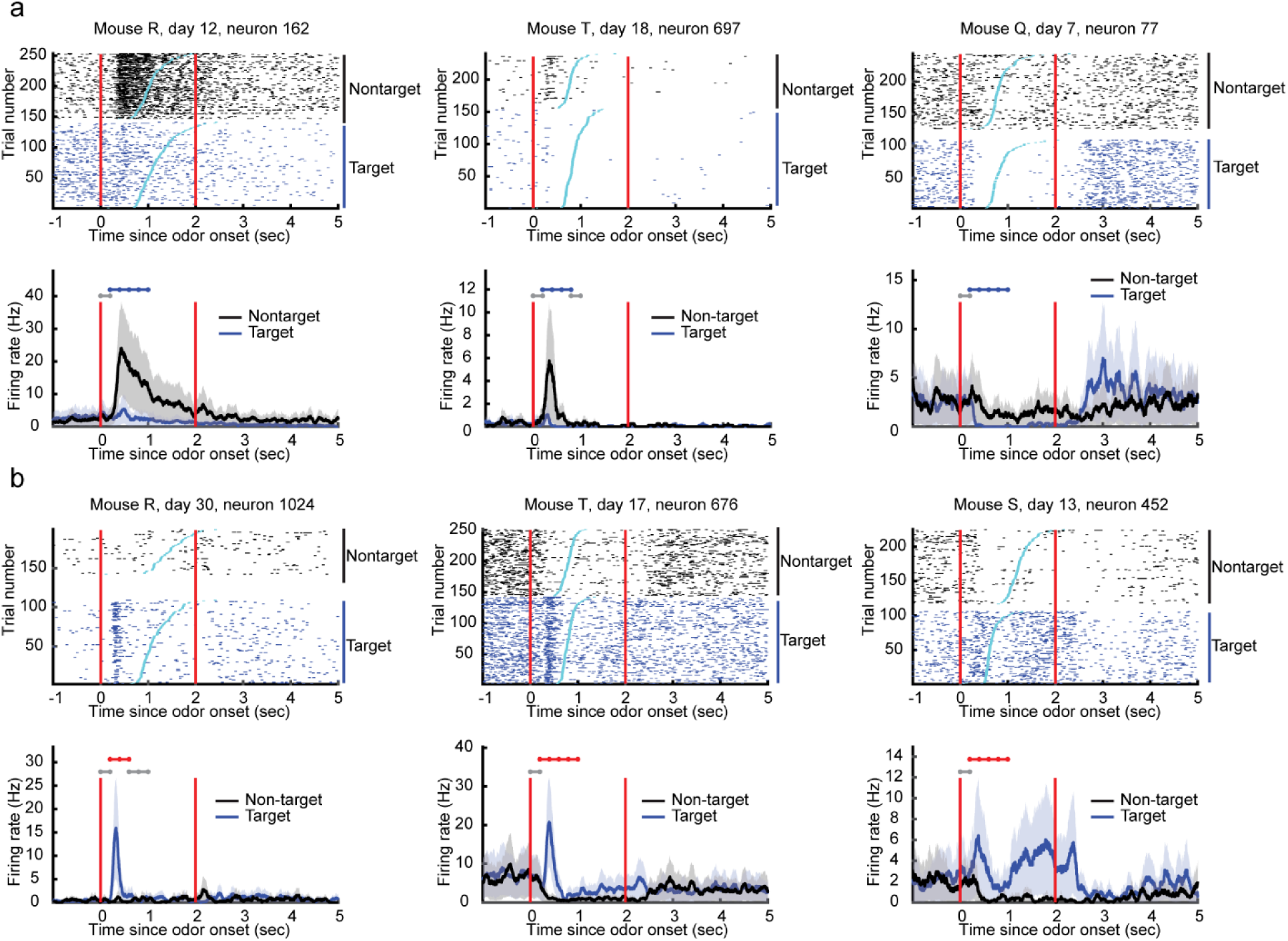
Some piriform neurons differentially modulate their firing rate to the target and nontarget odor mixtures. **(a)** Raster plots (top) and PSTHs (bottom) for three example neurons that respond significantly more to nontarget odor mixtures. **(b)** Similar plots for three other example neurons that respond significantly more to the target odor mixture.

**Figure 4.**
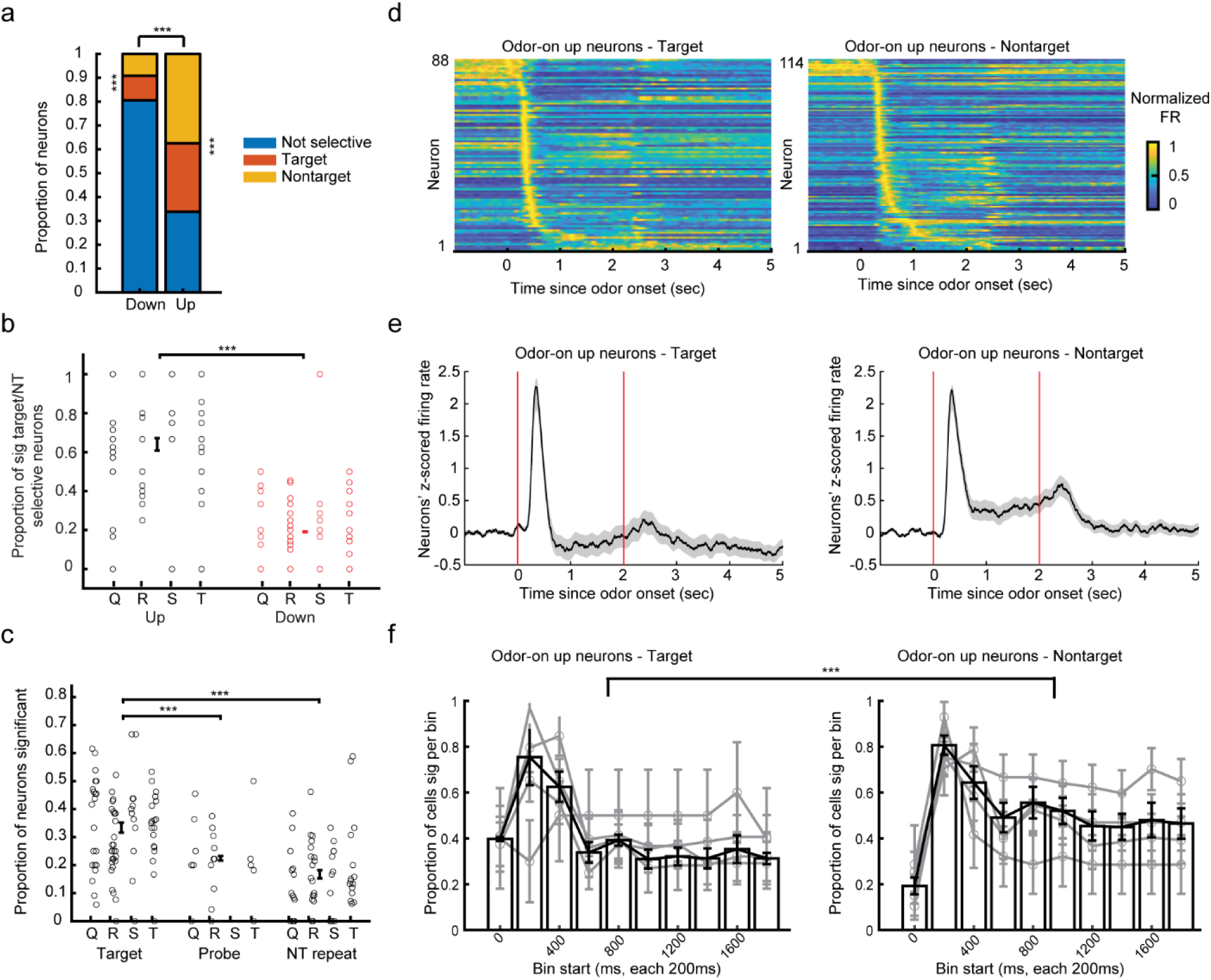
Piriform neurons respond differently to target and nontarget odors. **(a)** Proportion of all downwardly modulated neurons (left) and upwardly modulated neurons (right) that are significantly selective for target or nontarget odors. Binomial probability of up or down neurons being selective for target/nontarget is p < 7e-7 for both categories. A higher proportion of up-modulated neurons are selective for target/nontarget than down-modulated neurons (Chi-squared test p < 8.3e-35). Statistics details in Table 1, row 7. **(b)** Proportion of all upwardly modulated neurons (left) and downwardly modulated neurons (right) that significantly prefer target or nontarget odors across sessions of all four mice. 3-way ANOVA with direction (up or down), mouse, and session, details in Table 1, row 8. **(c)** The proportion of neurons that are significantly selective for target, probe, and nontarget repeats compared to nontarget odors across all sessions of all mice. Each small circle depicts a session. 3-way ANOVA with trial type, mouse, and session details in Table 1, row 9. Post-hoc Bonferroni comparison details in Table 1, row 10. **(d)** The normalized firing rate of upwardly modulated neurons that respond significantly more to target odors (left) and nontarget odors (right) **(e)** The mean baseline subtracted z-scored firing rates of upwardly modulated neurons that respond significantly more to target odors (left) and nontarget odors (right). Red lines depict odor onset and offset. **(f)** The proportion of upwardly modulated neurons that respond significantly more to target (left) or nontarget odors (right) that are significantly modulated in each 200ms bin. Gray lines represent the mean and S.E.M for each mouse separately. ANOVA details in Table 1, row 11. All error bars and shading represent mean +/− S.E.M. *** p < 3e-4.

These data show that a substantial number of piriform neurons discriminate between target and nontarget odor mixtures, more than other salient odor mixtures, and that the time course of a neuron’s response is related to its odor preference.

### Overlearning is characterized by latent improvements in piriform coding

After mice have learned the task, they exhibit consistent performance. Their accuracy as assessed by the choice index does not change across days (Figure 5a; Table 1, row 12). Additionally, the average lick latency does not change (Figure 5b; Table 1, row 13). We next wanted to see whether there was any covert learning that was not reflected in the mice’s behavior. We turned to see whether the responses of piriform neurons were changing during this same period.

**Figure 5.**
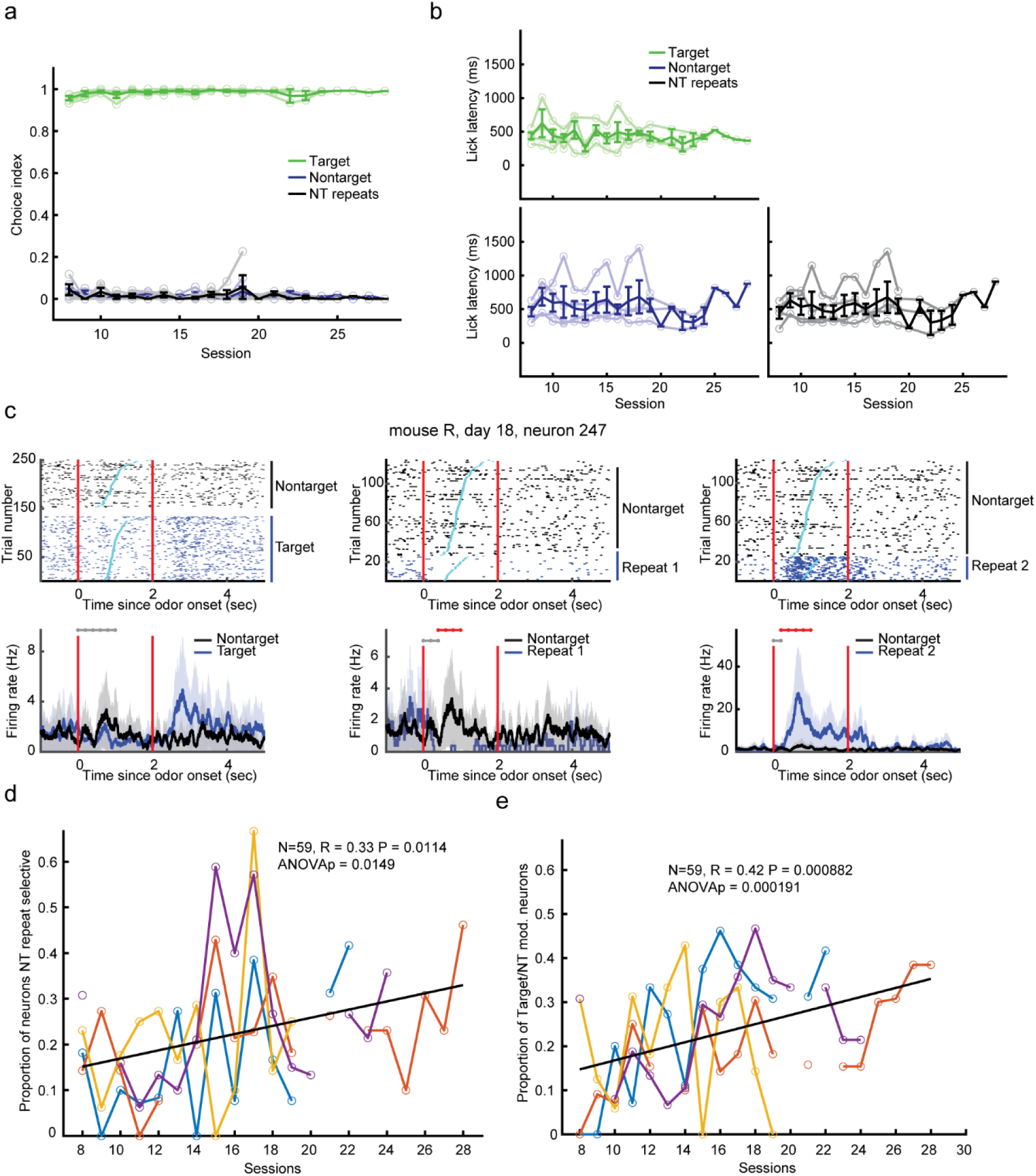
Target/nontarget task with nontarget repeats. **(a)** Choice index (1 for lick right, and 0 for lick left) for target, nontarget and the nontarget repeat trial types across overlearning sessions. No change in accuracy across sessions (ANOVA in Table 1, row 12). Dark colors are averaged across mice, light are individual mice. **(b)** Lick latencies (ms since odor-on) across sessions for each trial type. No change in latency across sessions (ANOVA in Table 1, row 13). Dark colors are averaged across mice, light are individual mice. **(c)** Raster plot (top) and PSTH (bottom) for an example neuron’s response to target compared to nontarget odors (left) as well as to two nontarget repeat trial types (middle and right). Note different y axis scale in bottom panels. **(d)** Proportion of neurons selective for either of the 1-2 nontarget-repeat trial-types vs nontarget trials across days. Black is a best-fit line. Each color is a different mouse. Significant change across sessions (ANOVA in Table 1, row 14). Note that breaks the data represent days that the mouse was trained but neural data was not available. **(e)** Proportion of neurons significantly selective between target/nontarget odors across days. Black is a best-fit line. Each color is a different mouse. Significant change across sessions (ANOVA in Table 1, row 15). All error bars and shading represent mean +/− S.E.M.

Our experimental design included two non-target odors that were repeated in each session (Figure 1b), which allowed us to interrogate how piriform cortex activity to these repeated non-targets evolved in the overlearning period. First, there are piriform neurons selective for a particular nontarget repeat, but they represented a smaller proportion than target-selective neurons (Figure 4c, 5c). The proportion of neurons on each day that were significantly selective for nontarget repeats (compared to all other nontarget odor mixtures; see methods) increases across days (Figure 5d; Table 1, row 14). Further, we found that the proportion of target selective neurons also increases across days (Figure 5e; Table 1, row 15).

To assess the possible consequences of this increase in the proportion of task-selective neurons, we performed categorical (target vs. everything else) decoding using the population of simultaneously recorded piriform neurons on each day. We performed 10-fold linear discriminant analysis decoding using the activity during the post-odor-on period of piriform neurons simultaneously recorded. We then compared this decoding accuracy to a shuffled distribution made by shuffling the labels of the category decision. We could readily decode category information for target, nontarget, and nontarget repeat trials (Figure 6a). Furthermore, although we were able to decode incorrect trials (average accuracy 60.0%, N = 1352, Monte-Carlo significance test P < 0.001), decoding was more accurate when the animal made the correct choice (Figure 6b insert; Table 1, row 16) and decoding was more accurate during sessions when the mouse performed better (Figure 6b). This is what we would expect if the mouse were using piriform cortex information to make its decision. We predicted that we would see an improvement in decoding across days, as the mice gain more experience, even though they were already at ceiling levels of performance. We found that the decoder performance increases across days for target, nontarget, and nontarget repeat trials (Figure 6c; Table 1, row 17-19). We also performed a boot-strap analysis to verify that this increase was significantly greater than expected given our data (p < 0.02), by shuffling the day of each session to generate a null distribution and found that the true slopes exceed this null distribution for each trial type (Figure 6d).

**Figure 6.**
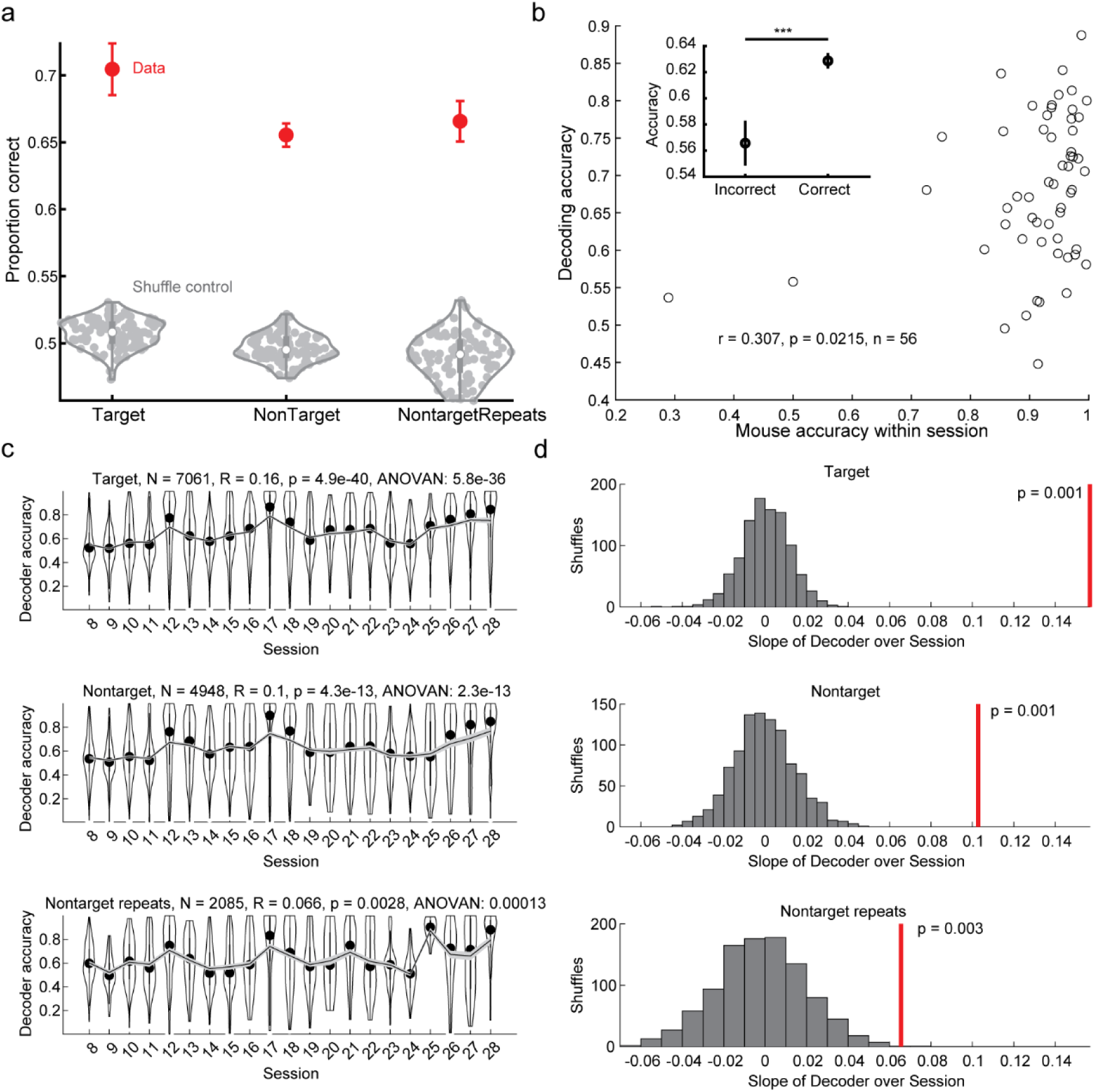
Decoding in the piriform population improves across sessions during the target/nontarget discrimination task. **(a)** Accuracy of decoding the categorical choice of the three trial types (red) compared to a shuffled distribution (gray clusters). All trials were averaged within each mouse. Error bars represent mean +/− S.E.M. across mice. Monte-Carlo significance test all P < 0.001. **(b)** The mean accuracy of category decoding plotted against the mouse’s average behavioral accuracy across all sessions. Inset: The mean accuracy of the category decoder for incorrect compared to correct trials after controlling for factors of repeat number, trial, session, and mouse. ANOVA details in Table 1, row 16. *** p < 0.005. **(c)** The mean decoder accuracy across days for each trial type. Top: target, Middle: nontarget, and Bottom: nontarget repeat. The relationship between session and accuracy assessed with a Pearson’s correlation (also significant in an ANOVA, details in Table 1, rows 17-19). **(d)** Pearson’s correlation calculated in c compared to a distribution of correlations calculated with shuffled (1000 times) session numbers. P-values from Monte-Carlo significance test.

Taken together, this data shows that during the overlearning period, while mice do not alter their behavior significantly, the underlying piriform population representation is changing. More piriform neurons are being recruited to be target and nontarget-repeat selective, and the discrimination of the two categories improves within the piriform population.

### The dissimilarity across category coding predicts accuracy on probe trials

Although the increase in task-selective neurons and the improvement in population decoding does not benefit the mice by improving their performance during over-training, we hypothesized that building more discriminable categories would benefit the mice when presented with more difficult decisions (Figure 7). After they were experts at the categorization task, three mice then moved on to sessions that included a new trial type, the probe trial. Probe trials consist of one of the three target odors and two randomly chosen odors from the nontarget odor pool (Figure 1a). Mice were able to perform well across all three types of probe trials (Figure 7a) and they improved across days of training with these trial types (Figure 1g, 7b; Table 1, row 20). We used pairwise similarity analysis to assess how strongly the piriform population separates target and nontarget representations (Figure 7c; Supplementary Figure 7; see methods: Population vector correlations). For each target or nontarget trial, we calculated the ‘target/nontarget similarity,’ which is the average population vector correlation between that trial (e.g. a target trial), and all of the opposite (e.g. nontarget trials) category’s population vectors (Figure 7d, each blue and green dot represents the ‘target/nontarget similarity’ for that trial and comes from averaging the blue and green lines). We predicted that the state of the target/nontarget representation during each probe trial could predict how the mouse would perform on that probe trial. To estimate the state of the cortex on each probe trial (pink dot in Figure 7d), we averaged the target/non-target similarity in the ten trials preceding the probe trial and plotted this value for each probe trial (Figure 7d-e; averaging the blue and green dots to get the pink dot in **d** and in **e**, averaging the black dots to get the blue and red markers).

**Figure 7.**
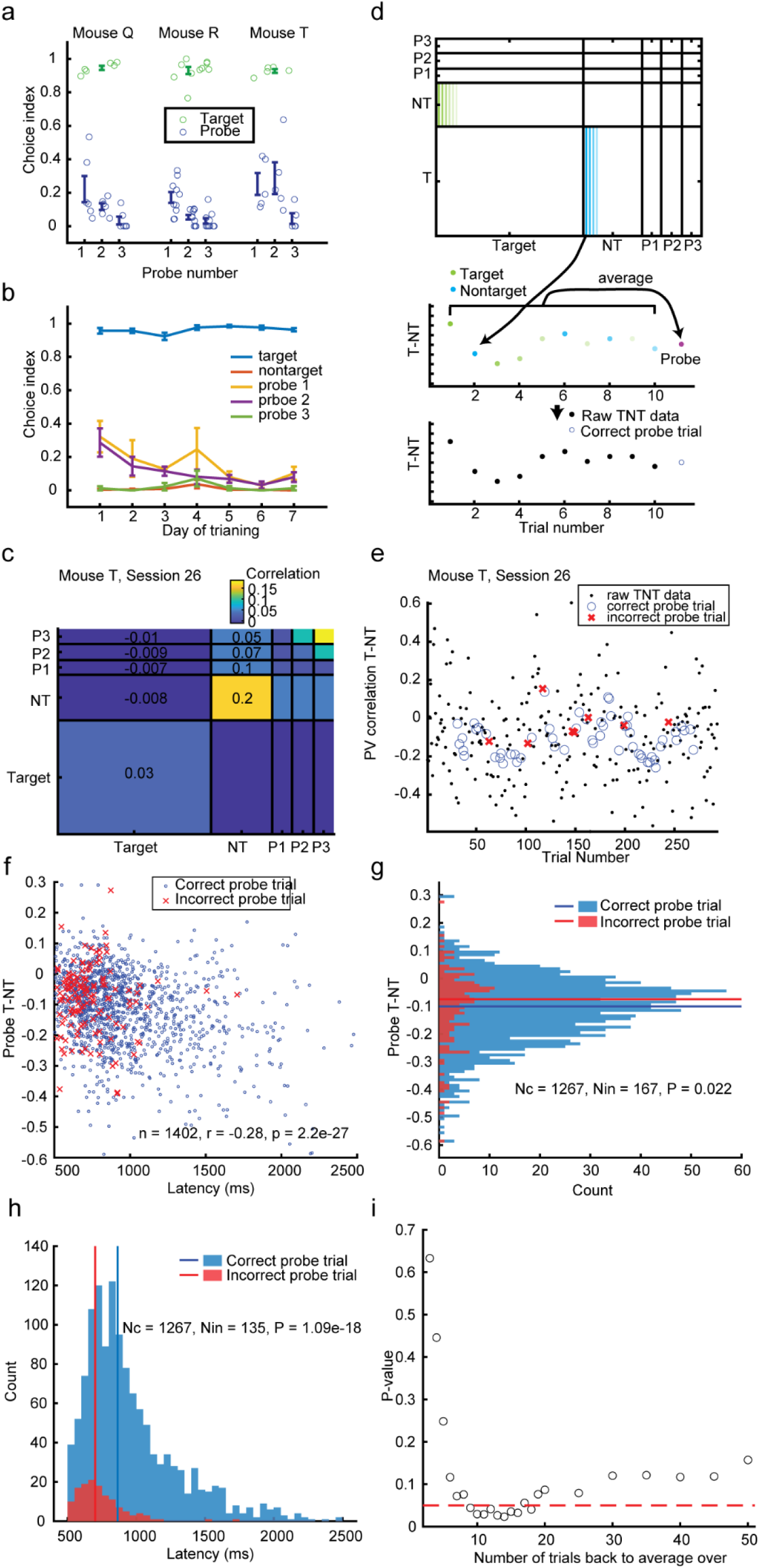
Dissimilar target/nontarget population responses relate to better subsequent performance on probe trials. **(a)** Choice index of three probe types across 3 mice. Each small circle is one session. **(b)** Choice index over days of training with the probe trials. Probe trials improve across sessions (see Table 1, row 20). **(c)** An example of population vector correlations across trials in one session, averaged within trial-type categories (P1-3 are probes 1-3). **(d)** Schematic of the procedure of determining the target-nontarget similarity (T-NT) just preceding the probe trials (probe T-NT). Top: The correlation between each target trial and all other nontarget trials was averaged to produce a T-NT for target trials (green), and vice versa for nontarget trials (blue). Middle: For probe trials (pink), the similarity is plotted as the average value of the 10 preceding trials, as a surrogate of the state of the piriform cortex when the probe trial arrives. The pink dot (probe trial) is the average of the blue and green dots (T-NTs for 10 preceding target and nontarget trials). Bottom: The example trial (here the mouse responded correctly) coded in the same colors as in **e**, which is an example session. **(e)** The T-NT for target, nontarget, and probe trials from the same example session as in **c** using the procedure described in **d**. The probe trial similarities are colored according to the mouse’s behavioral response – correctly classified as non-target (blue circles) or incorrected classified as target (red crosses). **(f)** The immediately preceding T-NT similarity for each probe trial plotted against lick latency for the probe trial across all mice (blue for probe trials with correct responses and red asters for incorrect responses). Negative TNT values represent a more differentiated piriform representation. There is a significant negative correlation (Pearson’s correlation R = −0.28, N = 1402, P = 2.2e-27). Monte-Carlo significance test p-value = 0.001. **(g)** The immediately preceding T-NT similarity of each correct (blue) and incorrect (red) probe trial. Correct trials have significantly less T-NT similarity than incorrect trials (One-sided signed-rank test, P = 0.022; Monte-Carlo significance test p-value = 0.013). **(h)** The lick latency of each correct (blue) and incorrect (red) probe trial. Correct trials have significantly higher latencies than incorrect trials (One-sided signed-rank test, P = 1.09e-18). **(i)** The same analysis computed in **g** but with a p-value computed across different look-back variables. Red dotted line depicts p = 0.05. Error bars represent mean +/− S.E.M.

Interestingly, probe trials with correct responses tended to have longer lick latencies than probe trials with incorrect responses (Figure 7h). This was also true for target and nontarget trials, but the speed accuracy trade-off was especially stark in probe trials (Supplementary Figure 8; also see Supplementary Figure 9 for more information on lick latencies).

Therefore, we looked at how this latency relates to the state of the piriform cortex. Across all sessions and mice, we found that the lick latency in a probe trial was inversely correlated with the target/nontarget representation similarity preceding a probe trial (Figure 7f; Pearson’s correlation R = −0.28, N = 1402, P = 2.2e-27). Negative similarity values represent a more differentiated piriform representation, and this differentiation tends to happen on trials that mice take more time to respond to. Furthermore, by looking at this same ‘target/nontarget similarity’ readout for probe trial types by whether they were correct (blue) or incorrect (red), we found that correct probe trials tended to have more dissimilar representations between target and nontarget in the preceding trials (Figure 7g). We then explored the parameter space of the number of trials we looked-back to interpolate the target/nontarget similarity and found that this relationship with accuracy is consistent and significant across look-back sizes of 9-18 trials (Figure 7i).

Therefore, although the improvement of the category representation in piriform during overtraining did not change behavior at the time, it was subsequently related to better performance in more challenging probe trials.

## Discussion

We recorded from pPC neurons as mice performed a categorization task to classify one target mixture of three odors as different from hundreds of other three-odor mixtures. Our main findings are (1) that the target odor mixture becomes over-represented with more pPC neurons distinguishing the target odor mixture than the other odors repeated in the task, (2) that even without any stark benefit in behavior, the pPC representation becomes more robust and selective with over-training, and (3) that this selectivity benefits behavior during a later perturbation with more difficult and ambiguous probe trails.

### Over-representation of relevant odors

A few types of learning occur that are reflected in an over-representation in individual piriform neurons’ selectivity. The largest over-representation was of the target odor mixture, likely showing the effect of both the repeated exposure of the odor-mixture and the reinforced dimension of the task (mice are rewarded for categorizing target and nontarget odor mixtures by licking to two different lick-ports). However, a significant proportion of pPC neurons are also selective for odor mixtures that mice were not forced to distinguish from others. A significant proportion of neurons differentiate probe trials, where only one of the three odors matches the target mixture, from nontarget mixtures, even while mice are rewarded for classifying these into the same category. Further, a significant proportion of neurons differentiate nontarget repeats (which are repeated more often in any given session), compared with other nontarget mixtures. This is the case while, again, mice are rewarded for classifying these trial types the same way as other nontarget mixtures. This proportion of neurons selective for the nontarget repeat also increases across sessions. These results are consistent with the interpretation in that there is a strong associative encoding within the piriform but also still strong pure sensory encoding as well. Instead of conforming completely to the task and collapsing all nontarget mixtures together, the piriform still can distinguish differences between these odor mixtures.

Although piriform cortex can still represent individual odor mixtures, its robust target coding means it already has all relevant information that can be used by a downstream area to compute a categorical decision. Our data does not speak to whether this learned over-representation is bottom-up, top-down, or produced from dynamics within piriform. However, it is likely that, while the afferent inputs are relatively set, the intracortical inputs are important for this learning (Bekkers and Suzuki, 2013; Bolding and Franks, 2018; Franks et al., 2011; Poo and Isaacson, 2011).

### Profile of Target-selective neurons’ activity

Our finding that target/nontarget selective neurons are more likely to be upwardly modulated is consistent with findings that inhibition is more broadly tuned than excitation (Bolding and Franks, 2017; Millman and Murthy, 2020; Otazu et al., 2015; Poo and Isaacson, 2009; Wang et al., 2020). In fact, the combination of fast upwardly modulated odor specific neurons and slower inhibited neurons could be indicative of circuit dynamics to affect population selectivity within the piriform (Bolding and Franks, 2018; Stern et al., 2018; Suzuki et al., 2022).

Further, we found that target-preferring neurons are more likely to increase their firing rates quickly after odor onset than nontarget-preferring neurons. Because there are hundreds of different combinations of nontarget mixtures, compared to the fixed three odors in the target mixture, these nontarget responses are likely drawing from a wider range of latencies earlier in odor processing (Carey et al., 2009; Haddad et al., 2013; Spors et al., 2006) and subsequently also have a wider distribution of feedback dynamics due solely to the huge distribution of odor mixtures it is based on. However, some of this effect could be due to learning as well, since recognizing the target odor-mixture (for example, by template matching) may be the fastest method of making an accurate decision using just identity coding. Perhaps this temporal refinement and truncation is due to recurrent circuitry within the piriform itself (Bolding and Franks, 2018; Bolding et al., 2020).

### Other notable neural correlates

We note that a number of lick-sensitive and odor-off sensitive neurons also exist in the piriform cortex. Lick correlates could arise from motor or taste responses (Maier et al., 2012). Although these were not the focus of our investigation, we reported the prevalence and were able to remove those neurons to concentrate on odor responses, and to assure ourselves that task decoding was not dependent on these responses. Odor-off responses have been reported in other species in olfactory receptor neurons and antennal lobe (Kurahashi et al., 1994; Nagel and Wilson, 2011; Nizampatnam et al., 2022), as well as in mouse piriform (Tantirigama et al., 2017). We expect these dynamics are likely due to feedback from within or outside the piriform circuit and not driven from the bulb, as we did not find a significant number of neurons that showed odor-off responses without any odor-on response.

### Neural changes despite behavioral stability

During a period of overtraining, when mice were already at ceiling in performance, we saw two types of changes in the pPC population. A greater proportion of neurons become target-selective, making the target coding an even more robust population response, reflected also in the increase in category decoding during this period as well. In addition, a greater proportion of neurons become selective for nontarget repeats, even as the task incentivized generalizing across all nontarget mixtures instead. This learning was purely due to exposure and did not sacrifice population decoding of the target/nontarget category decision, since the population decoding accuracy actually increases during that time for those nontarget repeat trials. Thus, when mice were experts and over-trained on this odor-discrimination task, latent learning of aspects of the task not currently relevant may have occurred without sacrificing task accuracy (Tolman, 1948). Improved discrimination of nontarget repeats from all other nontargets occurred through trials where the mouse was rewarded, but not to improve that discrimination. Thus, it could also be considered perceptual learning, perhaps occurring orthogonal to the reinforced dimension of target versus nontarget (Mandairon et al., 2006). This could be due to attention or the coincidence of reinforcement signals which arise to strengthen the task-relevant dimension, which is also theorized to support task-irrelevant perceptual learning in vision (Seitz and Watanabe, 2005).

The nontarget repeat mixture was also novel each day, meaning that this learning is a type of rule-learning different from stimulus-specific sensory learning (Slotnick, 2001; Slotnick et al., 1991; Wilson et al., 2004). Another implication of this finding is that as demonstrated with a simple decoder, any area downstream of the pPC can already have strong category information which could immediately be acted on without a second level of processing. This could mean that the mouse may be more reliant on the piriform than higher level areas such as the OFC during this type of task (Wang et al., 2020). Future experiments are required to know whether and when the piriform cortex and other areas are necessary for this task.

One caveat of this effect is that as recording sessions progressed, tetrodes were moved slightly by the experimenter each day to attempt to record from a new population of neurons. Although we cannot rule out layer differences, we do not think it is the most likely explanation for the effect. The overall trend of this movement was likely to be downwards (moving more ventrally), so earlier sessions of overtraining would be sampling deeper layers and later sessions sampling more superficial layers. As layer 2 is denser than the deeper layer 3 or more superficial layer 1, we calculated our neuron yield across these days as a proxy for density. We found that the yield decreases across days, which could correspond as a move from layer 2 towards 1 (2b to 2a), which would correspond to increasing proportion of semi-lunar cells. However, prior work has shown that semi-lunar cells are less selective for odors (Nagappan and Franks, 2021). Thus, this hypothetical is not consistent with the current knowledge of how layers and selectivity interact. Although we cannot fully rule this possibility out, we believe that due to this reasoning, along with the fact that tetrode movements were not large and can often lead to non-linear movements, the most reasonable explanation for the changing selectivity is still the continuous experience the mice accumulate.

### Importance of training history

Our data is consistent with prior work showing that rewarding animals for separation of odor types will lead to separation in piriform representation (Chapuis and Wilson, 2012; Shakhawat et al., 2014), but our data extends this from looking only within a mixture of odors to across different mixtures of odors. Interestingly, while Chapuis and Wilson’s experiments showed that experience reinforcing pattern separation leads to better pattern separation in pPC and experience reinforcing generalization leads to better generalization, we saw a slightly different pattern. Our task reinforced generalization across all the nontarget odors used for nontarget odor mixtures, and yet the pPC still showed pattern separation within that category, with nontarget repeats becoming more selective across experience.

The slight differences in population dynamics within these data highlight how important the details of an animal’s training history are to its cortical representation. In experiments in the olfactory bulb, training mice on easy or difficult odor discrimination leads to changes in robustness and efficiency in olfactory blub coding (Chu et al., 2017). The over-representation of the target odor mixture is a less efficient, but more robust coding scheme. It then becomes more and more robust with experience. One could imagine that with different protocol of training, a more efficient and less robust code may be produced instead. In fact, in other behavioral set-ups, pPC neurons have been shown to become less selective, making them better at categorization than aPC (Kadohisa and Wilson, 2006).

Although the changes in single neuron representation during the period of overtraining and the resulting increases in population decoding accuracy do not benefit the mice at the time, we hypothesized that they may benefit them when the task becomes more difficult. We found that in periods when over-trained mice had better separation between target and nontarget representations, they gave probe trials more time before responding, and were more accurate in their responses. This perturbation highlights the utility of piriform building a robust representation of the task. In an uncertain world where task difficulty and goals can change, the piriform is able to prepare for possible future reward without sacrificing current reward rate.

## Methods

### Surgery

C57BL/6J Mice from Jackson Laboratories, all male, 8-10 weeks old of age at start of the experiment. Experiments were conducted in accordance with Harvard University Animal Care Guidelines. Surgeries were performed on naive animals, and all behavioral training began after recovery from surgery. Mice were anesthetized (Ketamine/Xylazine 100 and 10 mg/kg, respectively). Following surgical implantation of a tetrode drive and a custom head plate, all mice we rehoused individually. A cranial window (~1 mm) was made over the dorsal skull at a location directly above the area targeted for electrophysiological recordings with the goal of implanting a tetrode bundle (eight tetrodes plus a 200-mm-diameter optic fiber to ensure stability). Posterior piriform coordinates: 0.5 mm posterior and 3.8 mm lateral from bregma, and 3.8 mm ventral from brain surface. To ensure stability of the head of the animal during behavior and recording at a later stage, a custom-made head plate (made of light-weight titanium; dimensions, 30×10×1 mm; weight, 0.8 g) was affixed to the skull. A shallow well was drilled over the posterior lateral skull, and a single skull screw was affixed at that location. A wire was attached to this skull for grounding electrophysiological recordings. In addition, a plastic cone was positioned around the tetrode drive, and capped with a removable lid, to prevent damage to the drive. Following the completion of the surgery, mice were given 1 week to recover before behavior training. Recordings were completed using openEphys (https://open-ephys.org/) for data acquisition and MClust (https://redishlab.umn.edu/mclust; https://github.com/adredish/MClust-Spike-Sorting-Toolbox) for spike sorting software.

### Behavior training

Mice were water restricted in compliance with approved protocols. Mice were acclimatized to the behavioral apparatus for one session in which they were allowed 30 minutes of free exploration with free water available. This was followed by an additional day in which they were head restrained and were allowed to lick for water from the two ports. In this session at any given time only one water port could deliver water and the mouse had to try both sides to discover which one. If the mouse collected the water drop from that port the next water drop had a 50/50 chance to be in either port, if the mouse only picked one side to lick, it never received additional drops unless it tried the other port. Generous manual delivery of water drops to the “correct” side occurred to help the mouse learn that there were two water ports and to avoid a side bias.

In the next phase of behavior training, the trial structure was introduced, where the odor was delivered for 2 seconds and a free drop of water was available at the correct lick port (for target the right lick port and nontargets the left lick port), and a ten second inter-trial-interval. This shaping phase of the behavior training lasted 1-2 sessions until the mouse refrained from licking during the inter-trial-interval.

The next training phase consisted of blocked trials – 3 of target and 3 of nontargets; in these trials the mouse had to lick the correct port to receive a water reward. When performance was greater than 70% correct for both target and nontarget trials, mice were moved to the next phase of training (typically 3-4 days of block sessions).

The next phase had no structure, but we forced trials on both lick ports. If the mouse started for example with a target and licked correctly, the next trial would be target or nontarget with equal probability; if the trial was incorrect, the next one would still be a target. When performance was greater than 70% correct for both target and nontarget trials, mice were moved to the next phase of training (typically 2 days of no structure force both sides).

The next phase had a random trial structure where every trial had a 50% chance of being target or nontarget. In the initial training phases if the mouse licked the incorrect port first but then made the correct lick during the decision period (odor on 2 seconds + 500ms), water reward was released from the correct lick port, however the trial was still counted as incorrect.

After performance was greater than 70% on both sides, an additional criterion was added wherein the lick choice had to be to the correct side only.

When behavior performance was greater than 90% for at least 3 days, the mouse was considered expert at the target/nontarget odor mixture task and probe trials were introduced in subsequent sessions.

### Data analysis

All analysis was performed using custom scripts in MATLAB R2022a, that are available on GitHub at https://github.com/ABernersLee/TargetPaper_20220822.

### Analysis of behavior

Odor onset is demarcated as when the last solenoid switched on. There was a consistent delay between this time and when the odor reached the mouse which was about a hundred milliseconds. We excluded the small number of trials in which the first lick came before 500 ms after odor onset. Choice index for target trials is the proportion of total trials that were correct, and for other trial types (nontarget trials, probes, nontarget repeats) it is 1 minus this value. The total number of trials was calculated in two different ways. In Figure 1 it was calculated during all trials, and trials when the mouse did not lick left or right to indicate a choice, were deemed incorrect. Only trailing trials where the mouse did not lick were truncated from the analysis (the last trial where the mouse licked was deemed the last trial). Supplementary Figure 1 was also calculated this way. In Figure 5, it was calculated only using trials in which the mouse made a decision, meaning that they licked left or right during the response period. The effects reported are stable between the two ways of calculating the choice index.

### Odor-on significance of individual neurons

We calculated the significance of firing rate changes due to odor presence or lick timing by comparing the firing rates during odor- or lick-related time periods in the task to the baseline firing rate (first 4 seconds of the trial and the last 4 seconds of the trial; 5 to 1 seconds before odor-on and 6 to 10 seconds after odor offset). To assess each time-bin’s significance we calculated the modulation [(bin-baseline)/(bin+baseline)] and compared it to a distribution of jittered spike trains where each trial was circularly shifted independently by a random amount of time. We calculated a p-value as the percentile of the absolute value of the real modulation compared to 1000 of those produced by a jitter procedure.

To identify significantly odor-modulated neurons we found neurons that had any of the five 200 ms bins in the 1 second after odor-on with a p-value of less than 0.01 (0.05/number of bins tested). Neurons were defined to be odor-on upwardly modulated if any of those significant bins were positively modulated, neurons with only negatively modulated significant bins were considered odor-on negatively modulated. Only neurons that were not found to be lick-modulated (see next paragraph) were designated as odor-modulated.

To find any neuron that could be contaminated by lick-modulation we aligned each neuron’s firing to either the odor (odor onset and odor offset) or the first lick (lick-on) and calculated the average peri-stimulus spike rate for both alignments in 200 ms bins. We evaluated the entire odor period (odor-on to odor-off; 2 seconds) and the lick-on adjacent period (1 second around the first lick; 400 ms before to 600 ms after). For comparing firing rates between lick-on and odor-on periods we only used trials where the lick occurred at least 800 ms after the odor-on time and only considered bins that had a p-value of less than 0.05. For neurons that had a positive odor-on modulation bin, if any lick-on adjacent bin was significant and had a higher firing rate than the firing rate in the positive modulation and significant bin, then the neuron was deemed lick-contaminated. For neurons that did not have a positive significant modulation bin, if any lick-on adjacent bin was significant and had a lower firing rate, then that neuron was considered lick-contaminated.

### Depicting the time course of neurons’ modulation

To show the responses of all the neurons in a given category (e.g. odor-on up or odor-on target preferring), we plotted a colormap of all the neurons across −1 to 5 seconds around odor onset. Each neuron’s smoothed firing rate was normalized by dividing by the maximum firing rate during that period. Then, to show the population response, we z-scored each neuron’s response during this same period and subtracted the average firing rate from the second before odor onset for each neuron.

### Significance of odor-identity modulation

To determine whether a neuron has a significantly different firing rate between two odor groups (target vs nontarget, nontarget repeats vs nontarget non-repeats, or probe vs nontarget) the difference in the firing rate in the same odor-on bins was used to identify odor-on significance (0-1 second in five 200 ms bins) between the two groups of trials. The unsigned average difference between the two groups was compared to the unsigned differences in a shuffled distribution where the group labels were shuffled 1000 times and the percentile of the real value compared to that distribution produced a p-value. A neuron was considered significantly selective for a group if any of the five bins had a p-value of less than 0.01 (0.05/ number of bins tested).

### Odor-off responses

To determine whether a neuron had a significant odor-off response, we used a similar method to that of determining odor-on significance but with a few more necessities to exclude responses that were not specific to the odor-off period. Neurons needed to satisfy three requirements: 1) Neurons needed to have at least one bin in the second after odor offset with a p-value less than 0.01 (0.05/five bins) compared with baseline. 2) Neurons needed to have a significant modulation (in the odor-off period) from the odor-on period. This was evaluated by taking the modulation in the 500 ms at the end of the odor-on period (1.5 to 2 seconds after odor-on) and comparing it to the 100 – 600 ms after odor offset. The modulation [(pre – post)/(pre + post)] was compared to a distribution of 1000 times that the trials were independently time-jittered and the same modulation was calculated. Neurons needed an unsigned modulation exceeding the 95th percentile of the jitter distribution (p-value of 0.05). 3) Neurons needed to have the same direction of modulation from both comparisons. For instance, if a neuron had a positive modulation compared to baseline periods, but a negative modulation compared to odor-on periods, this neuron would not be considered to have an odor-off response.

### Trial down-sampled comparisons

To check the validity of comparisons between proportions of target or probe/repeat selective neurons, which have different numbers of trials, we used a down-sampling procedure. Here, a reduced number of trials, two-thirds of the number of repeat or probe trials in a session (the trial-type with the least number of trials, thus equating the number of trials between the two types), was randomly chosen from the trial type (target, probe or repeat) and these trials were used to compare against the nontarget trials in that session in the same procedure as outlined above in *Significance of odor-identity modulation*. This procedure was performed 50 times for each neuron in each session, for each trial type. The mode of these 50 iterations’ p-values for each neuron was used to calculate the proportion of neurons that passed significance and these proportions were compared with a one-sided signed rank test.

### Changes during overlearning

To calculate the change in the proportion of selective neurons across days, we looked at two types of neurons. We looked at the proportion of nontarget repeat selective neurons and the number of target/nontarget selective neurons. We calculated a Pearson’s correlation and ANOVAs to determine changes in proportions across sessions.

### Population analysis

This section describes analyses applicable for both the decoding and population vector analyses. For each of the two experiments (probe and nontarget repeat) each session where this stimulus was included was analyzed separately (this includes one session where both experiments took place). For each simultaneously recorded set of neurons, the post-odor onset time window of 100 ms - 600 ms was taken for each neuron. We excluded neurons that fired fewer than five spikes summed across all trials in the post-odor onset time window. We excluded sessions with fewer than four neurons that passed this criterion. The trial by neuron matrix was z-scored for each neuron such that each neuron contributed equally to subsequent analysis.

### Decoding during overlearning

We performed 10-fold decoding (linear discriminant analysis) by randomly separating 90% of the trials into training data and 10% into test data. We did this 10 times and took the average of these 10 iterations. We used the classify.m function to perform linear discriminant analysis on the category decision (i.e. target vs everything else). To assess significance of the accuracy of this decoding, we performed 1000 shuffles whereby the category decisions were shuffled and the same procedure as described above was performed. We then performed a Monte-Carlo significance test to assess whether the accuracy of our decoding exceeded chance levels. To analyze changes across conditions (when the mouse was correct, or across days) we used the estimate of the posterior probability (using the third output of classify.m function) that the decoder chose the correct trial type as a more fine-grained measure of accuracy.

To analyze changes across the mouse’s accuracy, we performed an ANOVA with factors: repeat number, trial, day, mouse, and mouse’s accuracy. Both trial and day were continuous factors. We verified that there was a significant main effect of mouse’s accuracy in that ANOVA and reported the p-value. We then used multcompare.m to assess the direction of the difference. We reported the Tukey’s post-hoc test p-value and plotted the estimates and standard errors of each group.

To analyze changes across days, for each trial type separately (e.g. target) we performed an ANOVA with factors: repeat (when applicable), trial, day, mouse, and mouse’s accuracy. Both trial and day were continuous factors. We verified that there was a significant main effect of day and reported the p-value.

We then performed a Pearson’s correlation of the accuracy across days and reported the correlation and significance values. Finally, we generated 1000 distributions of correlation values with shuffled day-labels, in order to assess whether the changes we saw across days was significant given our data. We assessed this using a Monte-Carlo significance test and reported the p-value.

We also performed this same analysis, except using the residuals from a fitted model using all factors except day (not shown). Specifically, we took the marginal mean category decoder accuracy from an ANOVA with factors repeat number, trial, session, and mouse with only correct trials. We then performed the same analyses as stated above (Pearson’s correlations and Monte-Carlo significance tests on shuffled distributions) to verify that these results were also significant.

All of these analyses were also repeated excluding lick-contaminated neurons and excluding trials where a lick comes before 600ms post-odor onset and the results were similar and similarly significant.

### Population vector correlations

To generate population vector correlations, a form of representational similarity analysis, correlations between population vectors for each trial was computed using the corrcoef.m function. These correlation coefficients were then z-scored. To visualize within and across group differences, trials were averaged within trial-types (e.g. probe vs. target or probe vs. all other probe trials) while never including a trial’s correlation with itself.

#### Interpolating target/nontarget similarity to probe trials

To calculate the target-nontarget population (T-NT) similarity on a given trial, we took the population vector correlations between each target trial and all nontarget trials, and vice versa. Then, for each probe trial, we calculated the Probe T-NT: an average of the 10 (or another look-back window) prior trial’s T-NT similarities and assigned that as the probe trial’s value.

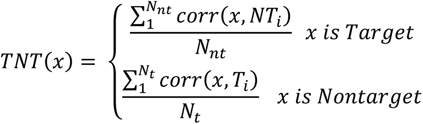

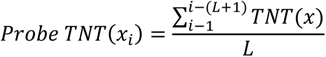

Where *L* is the number of trials to look back (10 in main analyses), *T* is a target trial, *NT* is a nontarget trial, *N_t_* is the number of target trials, *N_nt_* is the number of nontarget trials, and *i* is the trial number.

We compared the T-NT similarities for probe trials with those probe trials’ lick latencies using a Pearson’s correlation. We also performed a non-parametric significance test of that Pearson’s correlation by generating a distribution of 1000 correlations each using a different shuffled indices of latencies and T-NT similarities. We then tested whether this correlation is less than expected given the shuffled distribution by calculating the Monte-Carlo significance test.

Next, we compared these values in probe trials that were correct vs incorrect and in long or short latency probe trials using one-sided signed-rank tests. We also performed a performed a non-parametric significance test by taking the difference in the medians (correct-incorrect) and computing this 1000 times shuffling the accuracy values. We tested whether this difference was less than you would expect given the shuffled distribution by calculating the Monte-Carlo significance test. For the correct vs. incorrect comparison, we also repeated these analyses with look-back windows of 3-50 trials to look back and average over and found consistent significant results with a trial history of 9-16 trials.

### ANOVAs

All ANOVAs were performed using anovan.m (if more than one factor) or anova1.m functions. All trial or session factors were considered ‘continuous’. To look at the residual changes across one factor of interest (for instance across sessions) after accounting for the other factors in a model, we looked at the p-value for the factor of interest in a full model. Then we used the residuals from a fitted model using all factors except the one of interest, in order to visualize and perform post-hoc statistics to describe the trend (e.g. Pearson’s correlation).

## Acknowledgement

We thank Vikrant Kapoor for help with the experimental set up, and Kenneth Blum and Gautam Reddy for feedback on the manuscript. This work was supported in part by the NIH (R01DC016289).

## Supplementary Figures

**Supplementary Figure 1.**
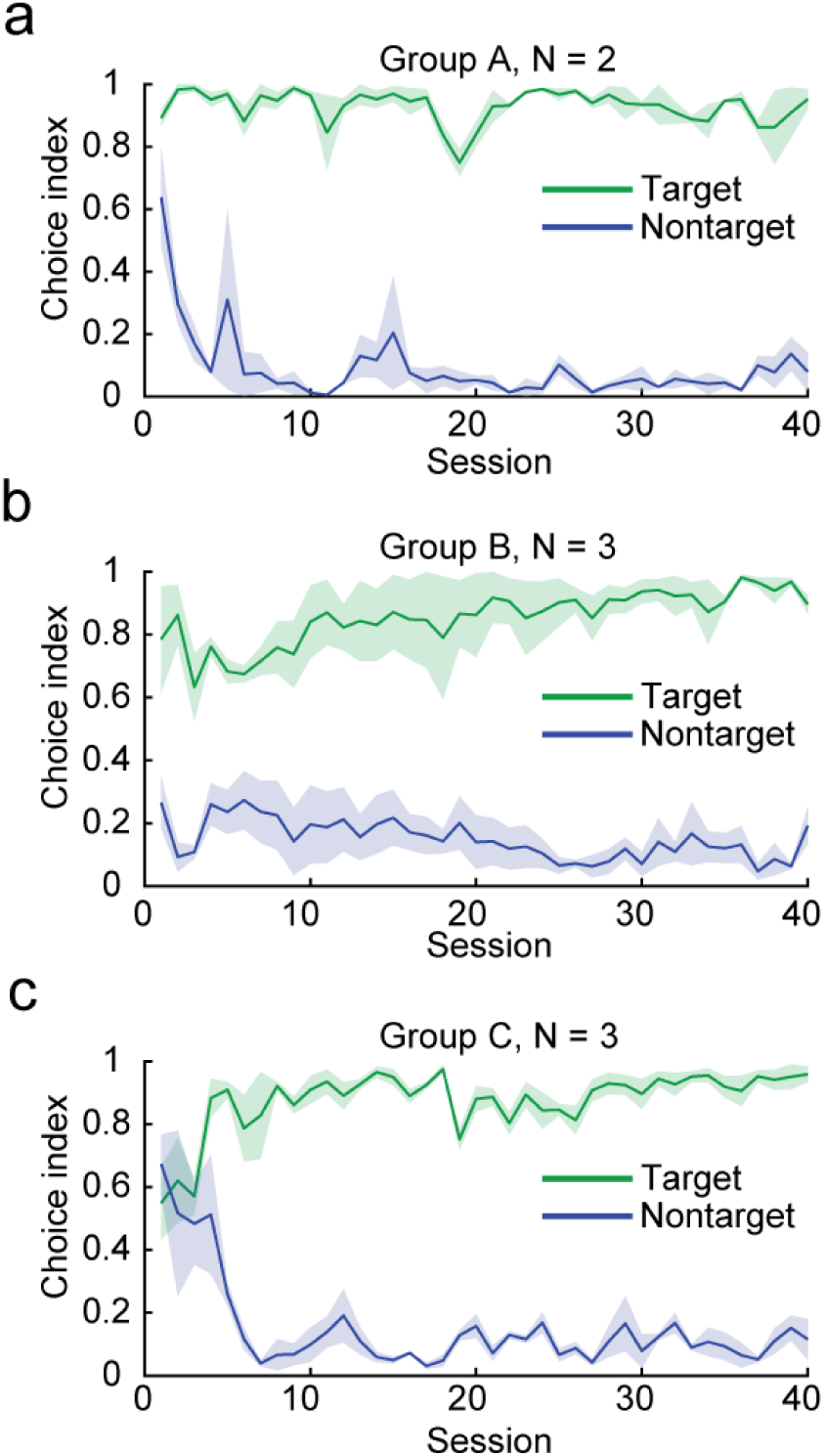
Different target odor mixtures – related to figure 1. Average choice index across the first 40 sessions for groups of mice trained on different target mixtures. Session 1 for each mouse is the first session with blocked trials. **(a)** Mouse group A trained on target mixture: Heptanal, Propyl Acetate, Isoamyl acetate. **(b)** Mouse group B trained on target mixture: Allyl buterate, Ethyl valerate, Methyl tiglate. **(c)** Mouse group C trained on target mixture: Ethyl tiglate, Allyl tiglate, Methyl tiglate. Shading represents mean +/− S.E.M.

**Supplementary Figure 2.**
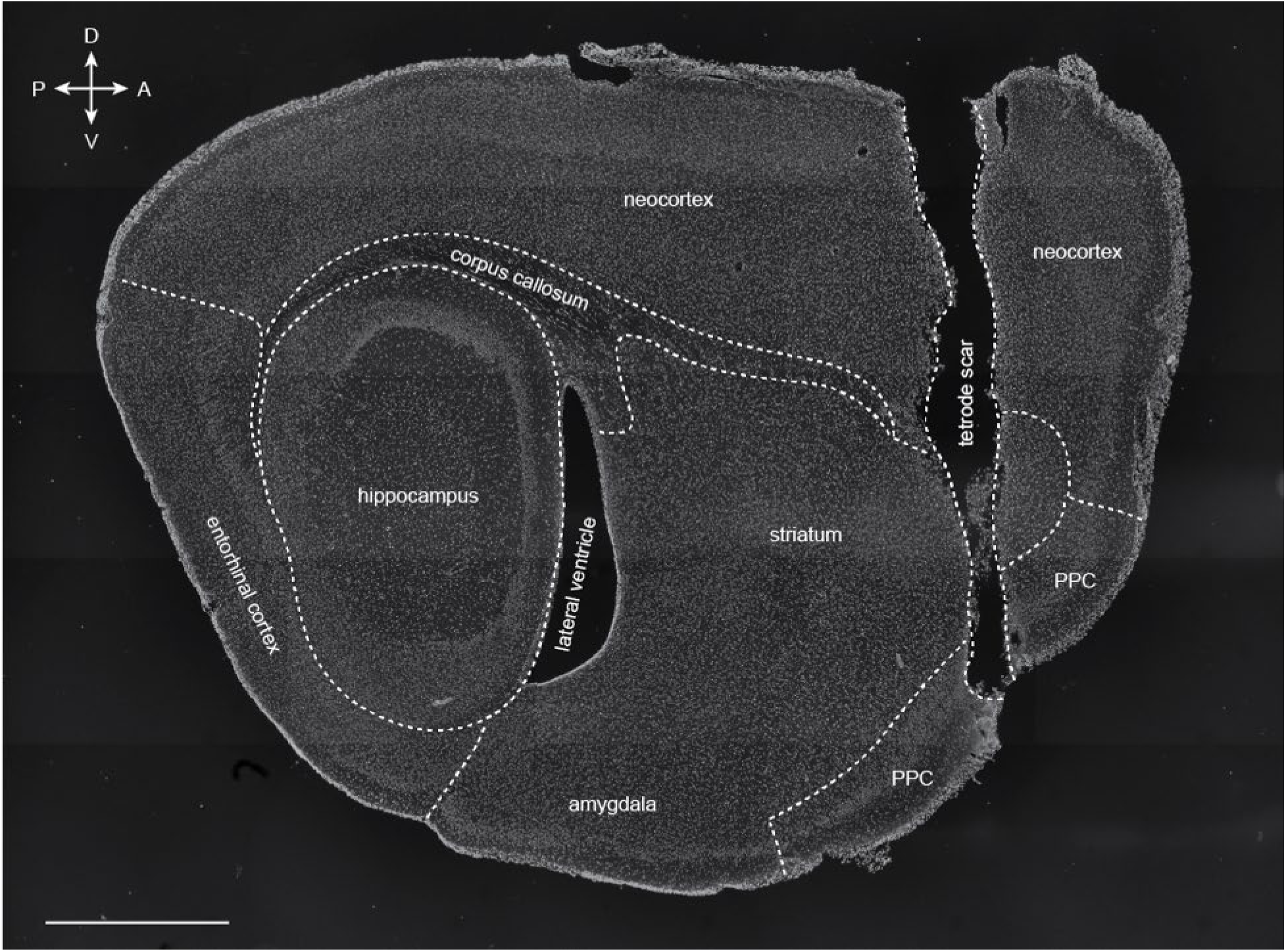
Post-mortem confirmation of tetrode placement – related to figure 1. This sagittal section comes from the right hemisphere of mouse S. The oval shape of the hippocampus, the shape and position of the lateral ventricle, as well as the absence of lateral olfactory tract confirm that the tetrodes were placed in the PPC and not the APC. A: anterior. P: Posterior. D: dorsal. V: ventral. Scale bar: 1mm. DAPI staining. We did not trace the border between striatum and amygdala as we could not confidently determine it.

**Supplementary Figure 3.**
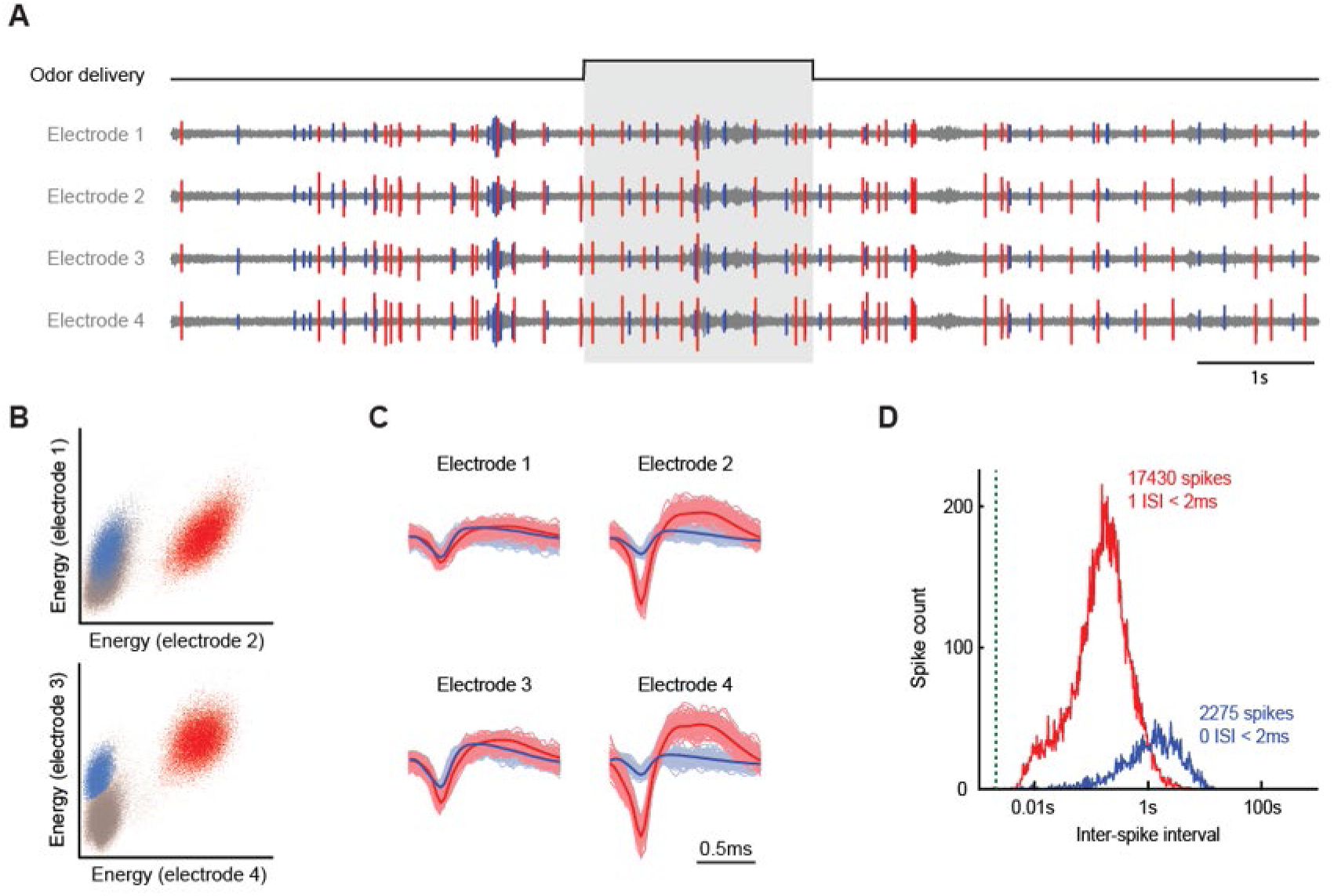
Example of spike sporting – related to figure 1. **(a)** Example of tetrode recoding. The four electrodes are from the same tetrode. Two single units are shown in red and blue. Grey: odor delivery. Example from mouse E, session 2-22-2017, tetrode #4. **(b)** Single unit clustering. The red and blue units are the same as (a). **(c)** Clustered single units. The red and blue units are the same as (a). The lighter traces show the 100 first extracellular action potential of each unit superimposed. The darker traces show the average extracellular action potential. **(d)** Single unit confirmation. The red and blue traces belong to the red and blue units from (a). The inter-spike interval analysis reveals the presence of a refractory period.

**Supplementary Figure 4.**
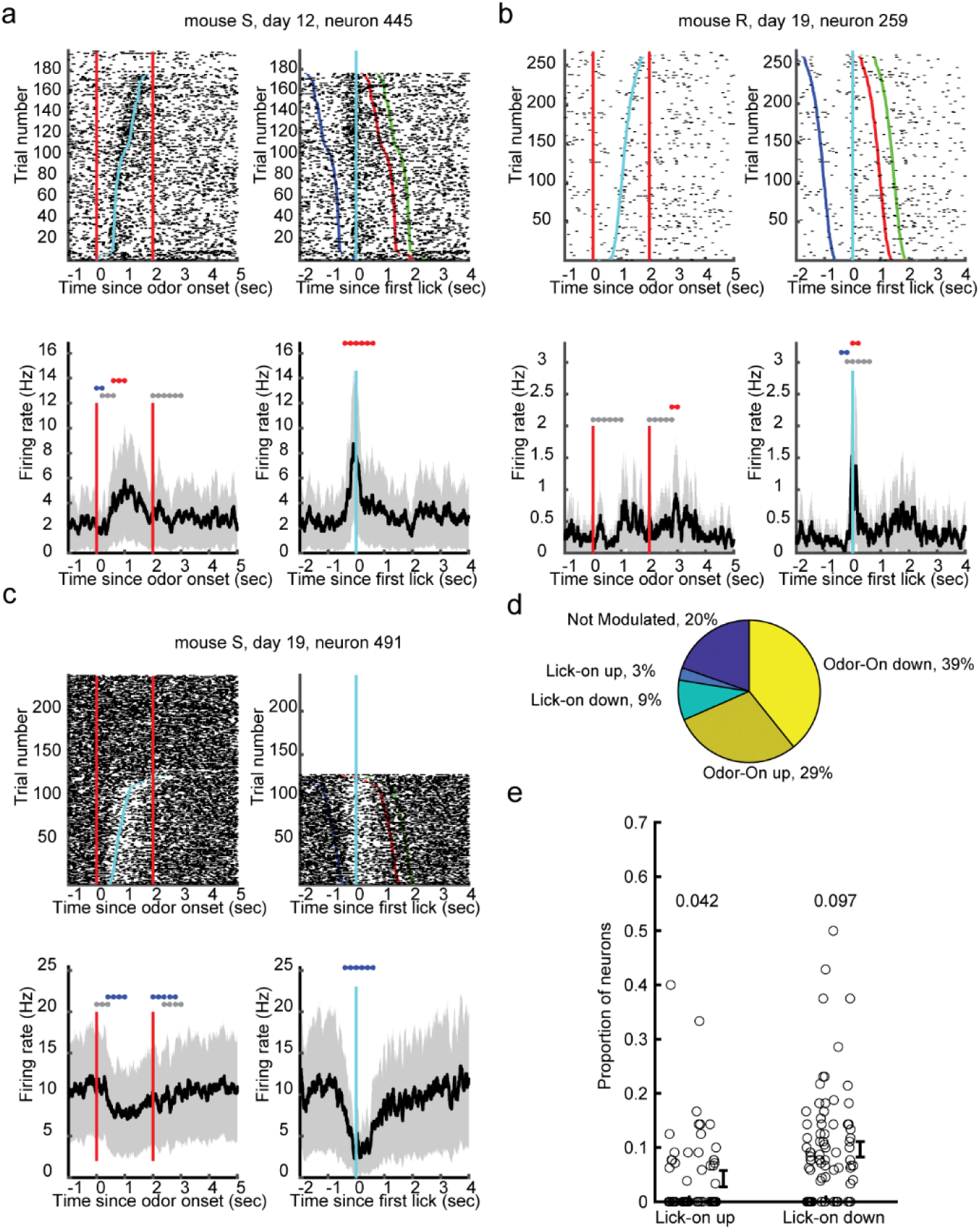
Lick-responsive neurons – related to figure 2. **(a-c)** Three example neurons that are modulated at first lick or reward. Raster plots (top) and PSTHs (bottom) of odor aligned (left) or lick-aligned (right) trials. Red lines depict odor onset and offset, blue dots depict lick times. Green dots represent the end of the response period. **(d)** Proportion of the total number of neurons recorded with significant task-relevant responses. **(e)** Proportion of neurons in each session (circles) with lick-on up and lick-on down responses. Average proportions are above each group. Error bars and shading represent mean +/− S.E.M.

**Supplementary Figure 5.**
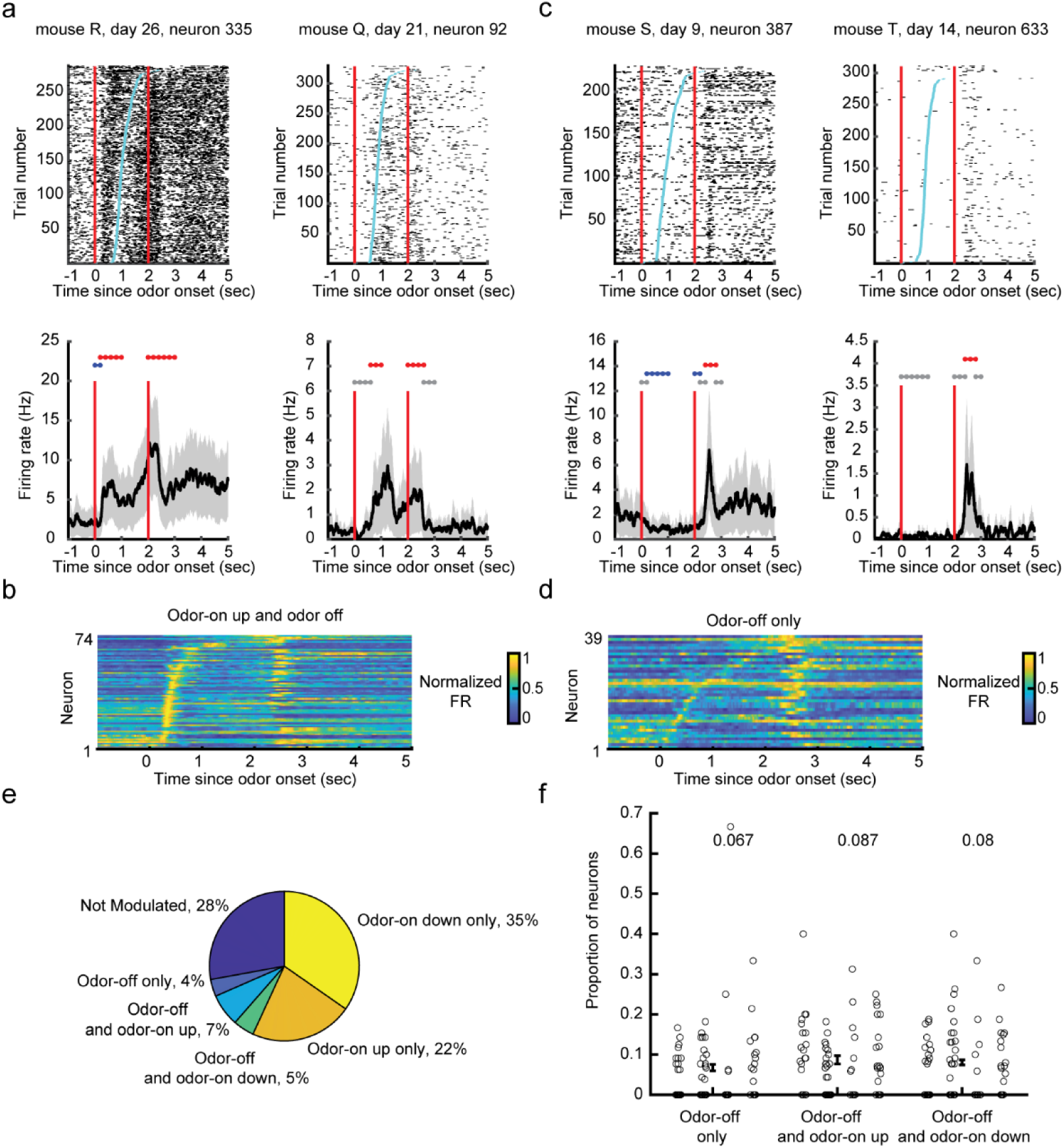
Odor offset responsive neurons – related to figure 2. **(a)** Two example neurons (left and right) that have both a significant odor-on up and odor-off response. Raster plots (top) and PSTHs (bottom) of odor aligned trials. Red lines depict odor onset and offset, blue dots depict lick times. **(b)** The normalized firing rate of all odor-on up and odor-off responsive neurons. **(c-d)** Same as in **a-b** but with odor-off only neurons. **(e)** Proportion of the total number of neurons recorded with significant task-relevant responses. **(f)** Proportion of neurons in each session (circles) with different odor-off responses. Average proportions are above each group. Error bars and shading represent mean +/− S.E.M.

**Supplementary Figure 6.**
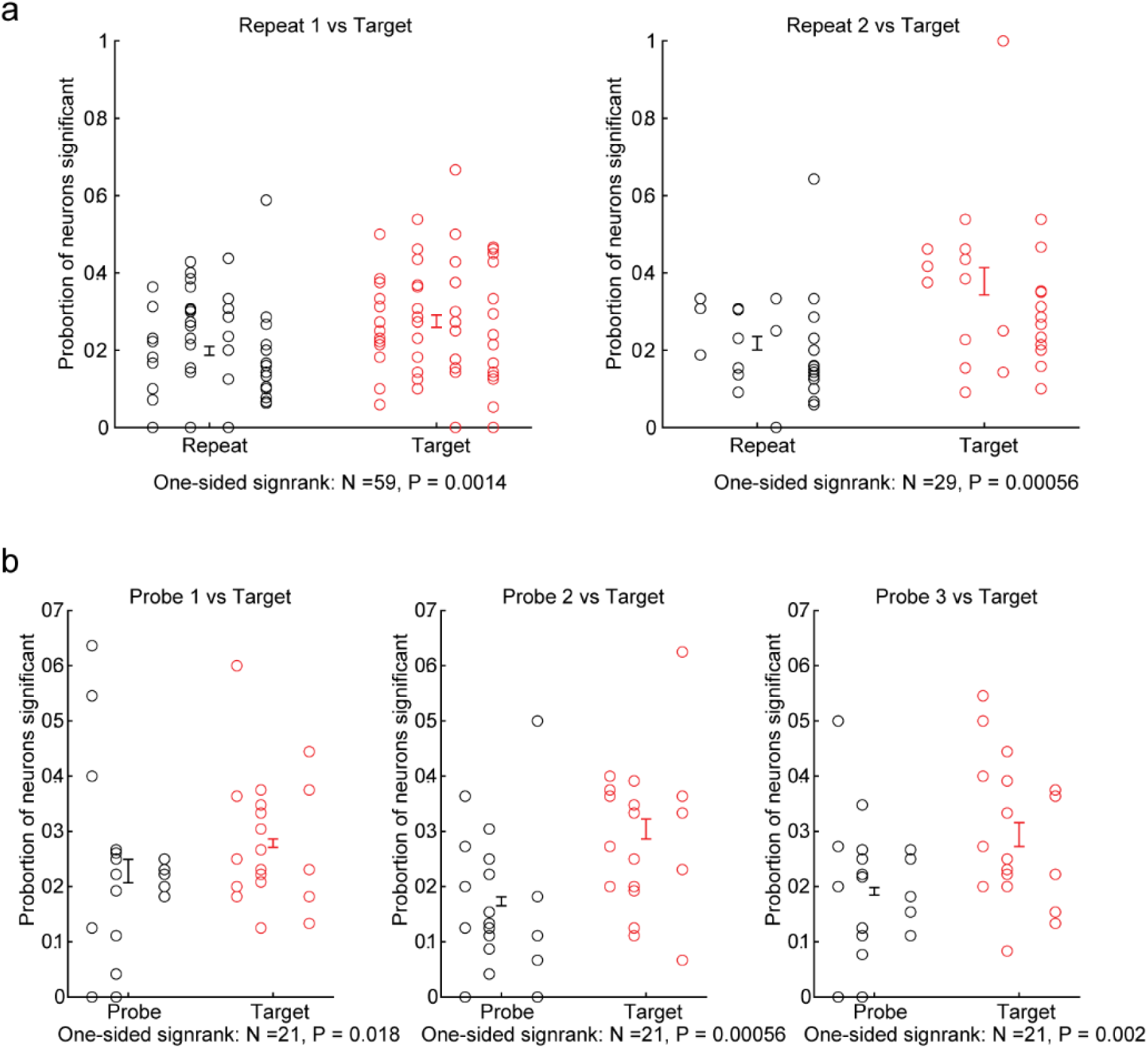
Paired down-sample control – related to figure 4. We repeated comparisons while down-sampling the number of trials to control for the uneven number of trials across trial types. Details of this procedure are in the *Trial down-sampled comparisons* section of the methods. **(a)** Proportion of neurons significantly selective for NT repeat (black) and target (red) trials. One-sided signed rank test to determine whether there is a higher proportion of target selective neurons than NT repeat selective neurons separately for each of the two possible repeats within a session (left vs. right). **(b)** Proportion of neurons significantly selective for probe (black) and target (red) trials. One-sided signed rank test to determine whether there is a higher proportion of target selective neurons than probe selective neurons separately for each of the two possible repeats within a session (left vs. right). See methods for more details. Error bars represent mean +/− S.E.M.

**Supplementary Figure 7.**
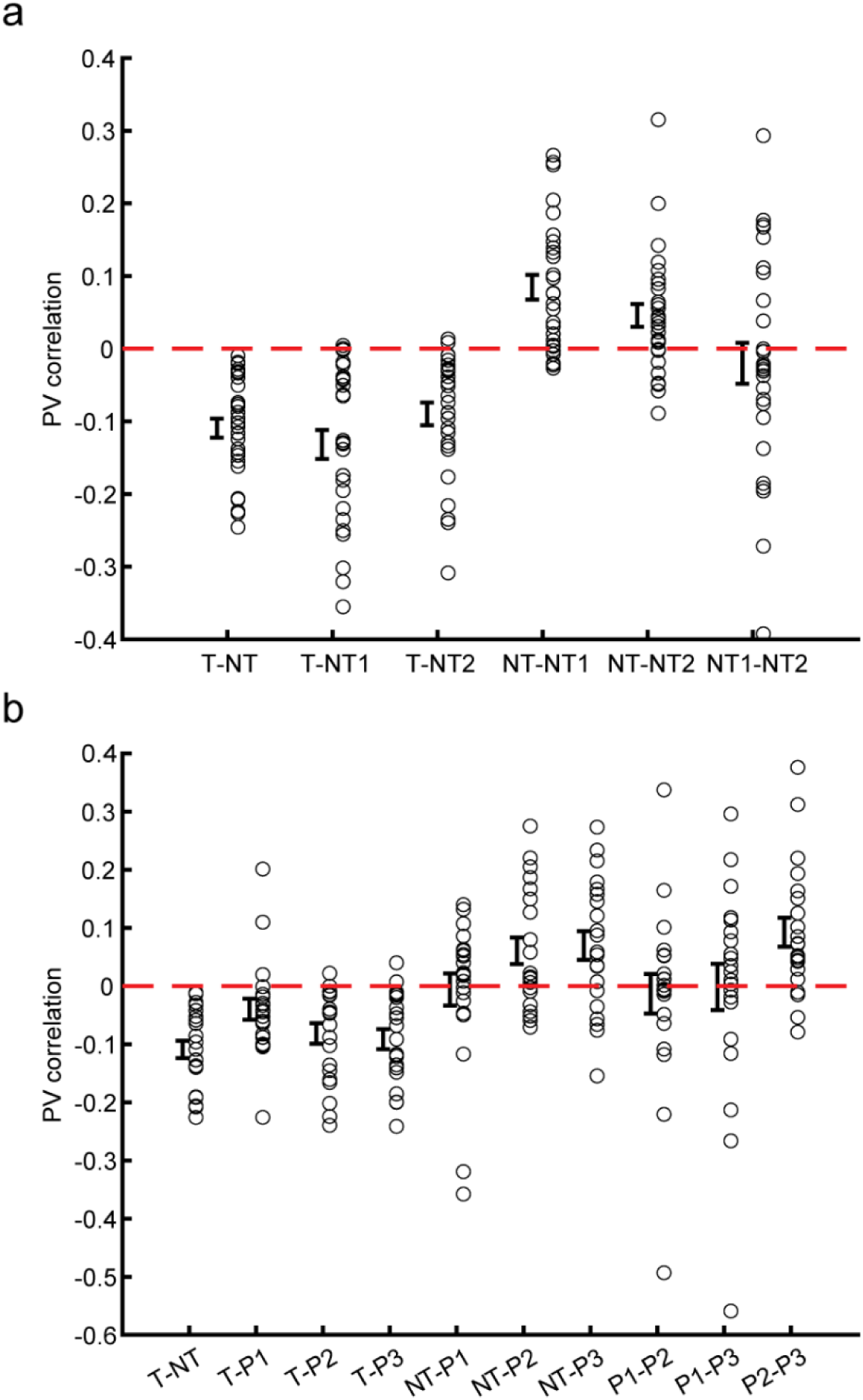
Average population vector corrections across groups - related to figure 7. Average population vector corrections for each session (open circles) between each trial type category in **(a)** over-learning sessions with at least two nontarget repeats, and **(b)** probe sessions. T – Target, NT – Nontarget, NT1-2 – Nontarget repeats 1-2, P1-3 – probe 1-3. Error bars represent mean +/− S.E.M.

**Supplementary Figure 8.**
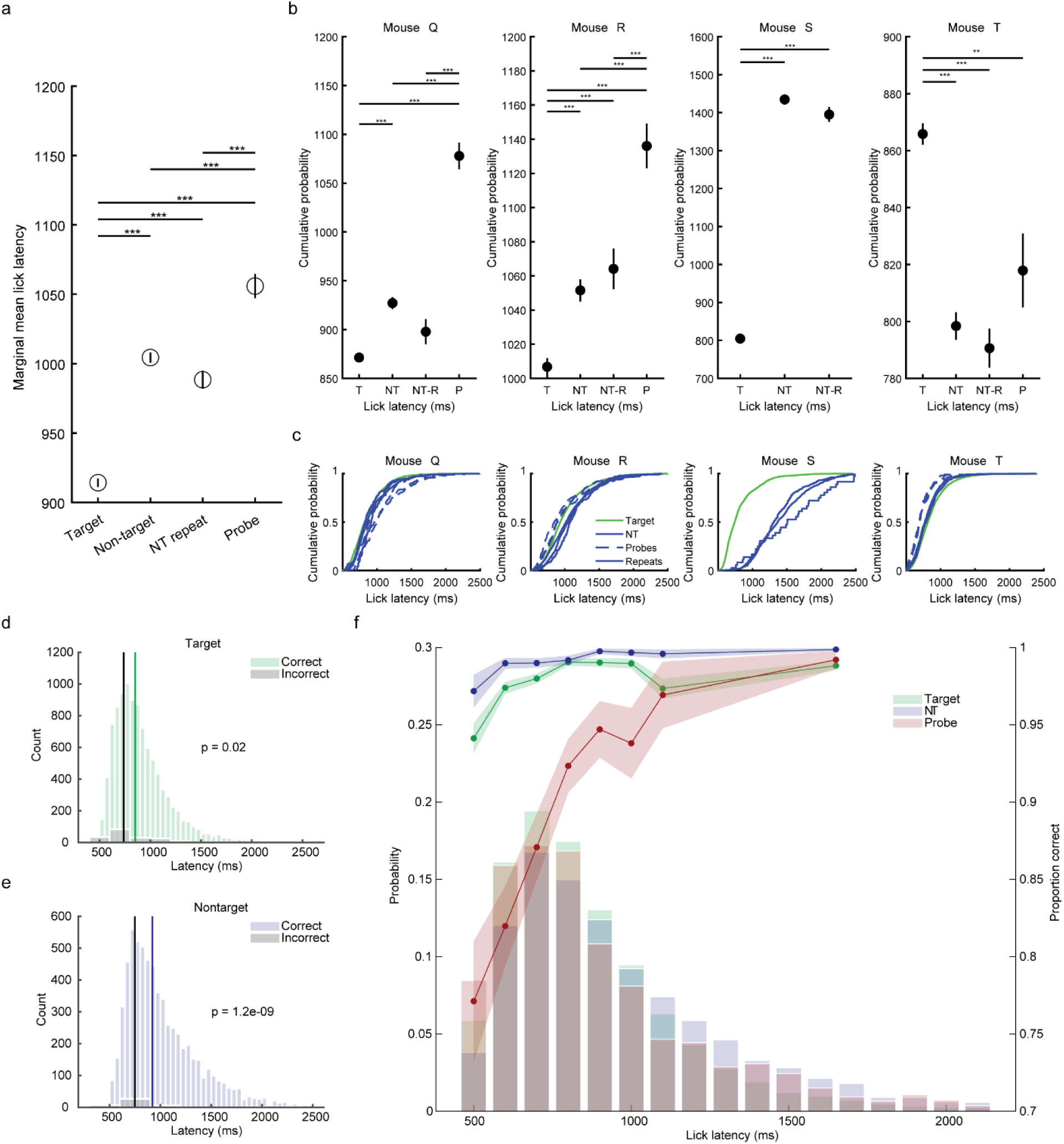
Lick latency across trial types – related to figure 7. **(a)** Trial-type lick latency differences. Marginal mean lick latency by trial type from a 3-way ANOVA with trial type, mouse and session. Main effect of trial type: F(20288,3) = 176.3, P = 7.2e-133. Main effect of mouse F(20288,3) = 853.9, P = 0. Main effect of session F(20288,3) = 973.2, P = 9.6e-209. Post-hoc Bonferroni tests between trial types shown with asterisks **(b)** Marginal mean lick latency for each mouse by trial type from a 2-way ANOVA with trial type and session. All mice showed a main effect of trial type (P < 1e-20 in all mice). Post-hoc Bonferroni tests between trial types shown with asterisks. * P < 0.05, ** P < 0.01, ** P < 0.001. Error bars in a represent mean +/− S.E. Note different y axis scales. **(c)** Lick latency distributions for all trials of target, nontarget, nontarget repeats, and probe trials, for each of the four mice. **(d)** Lick latencies for all target trials and **(e)** nontarget trials, separately for correct and incorrect trials. Only trials during probe sessions were used. **(f)** Left axis: A histogram of lick latencies during probe sessions by trial type. 100 ms bins. Right axis: The proportion of correct trials as a function of lick latency. Only probe sessions were used. The last data point includes latencies of 1200 – 2200 ms. Shading represents mean +/− S.E.M. We only considered trials in which the first lick came more than 500 ms after odor onset.

**Supplementary Figure 9.**
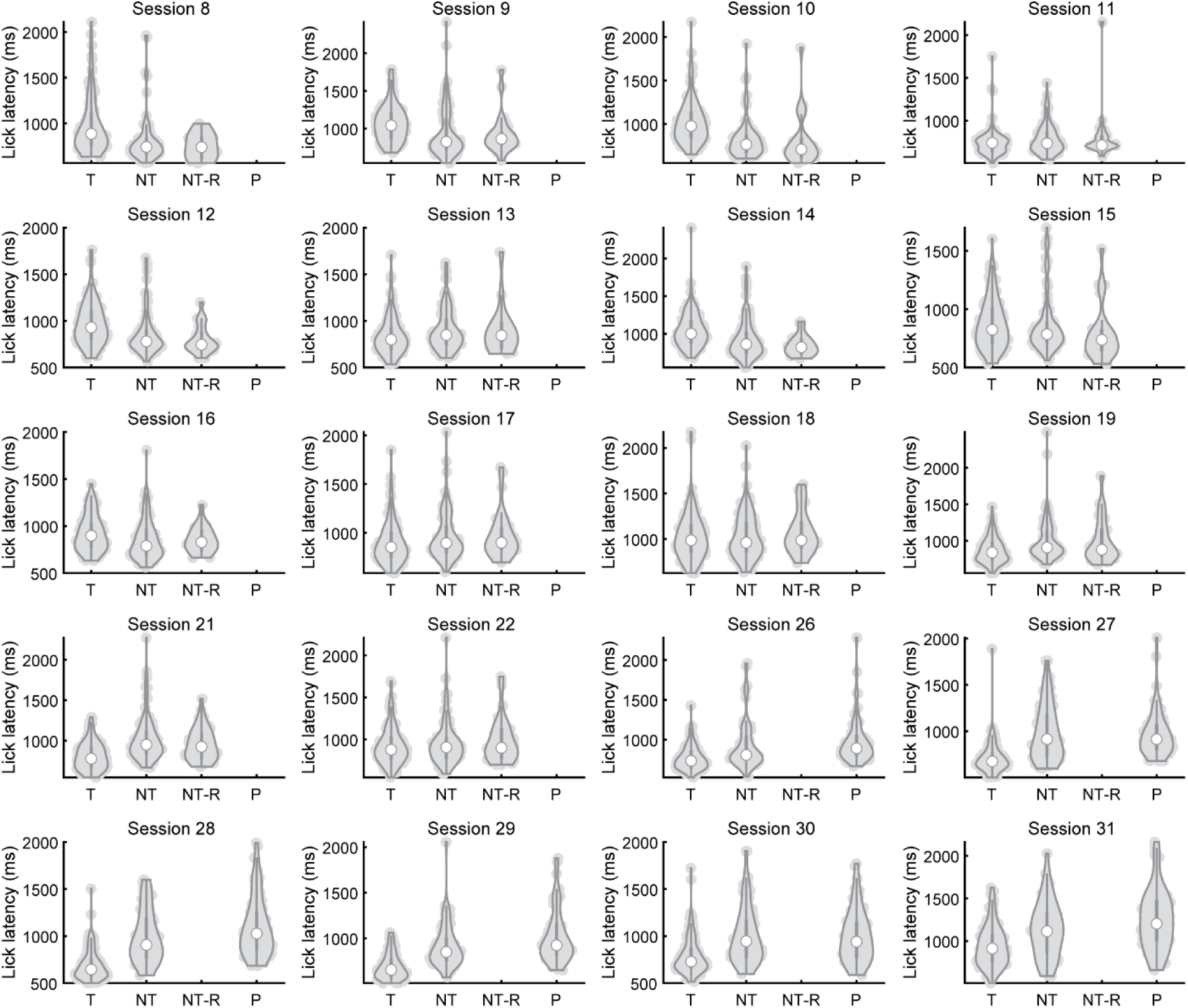
Variability in raw lick latency across sessions–related to Figure 7. Violin plots of mouse Q’s lick latencies in each session. X-ticks denote trial types. T: target; NT: nontarget; NT-R: nontarget repeat; P: probe.

